# Arginine activates a conserved sphingolipid defense axis against infection

**DOI:** 10.64898/2026.11.03.763873

**Authors:** Shuncai Han, Mengge Li, Geoff M. Gurr, Liette Vasseur, Minsheng You, Xiaofeng Xia

## Abstract

Metabolism is generally viewed as supporting immune responses rather than acting as a direct mediator of host defense. Here, we show that arginine directly activates the sphingolipid metabolic pathway in *Plutella xylostella*, promoting ceramide accumulation and consequently exerting antibacterial effects. Furthermore, we find that arginine synthesized by the host gut bacteria can activate this pathway, revealing an independent metabolic defense mechanism driven by gut bacterial arginine, which operates independently of classical immune cascades. Cross species validation shows that the ceramide biosynthesis pathway is highly conserved across eukaryotes, and its antipathogen effects display potential universality across insect species. This study opens new directions in gut innate immunity research, provides novel targets for the development of antimicrobial agents against insect pests, offers important reference value for eukaryotic pathology studies, and provides new insights into the development of novel therapeutic strategies against drug resistant bacterial infections.

## Introduction

For more than a century, studies of immunity have primarily focused on how host proteins recognize pathogens and activate downstream defense programs [1–4]. Although metabolism is increasingly recognized as an integral component of immune responses, it is generally regarded as supporting rather than executing host defense [5–8]. Whether metabolic pathways can function as autonomous immune systems, independent of canonical signaling cascades, remains a question that warrants systematic exploration.

Increasing evidence suggests that metabolites can serve as direct effectors of host defense rather than merely supporting immune responses. For example, itaconate accumulates in activated macrophages and suppresses bacterial growth through inhibition of isocitrate lyase [9–11]. Sphingolipid metabolism represents another promising candidate for metabolic immunity. Its central metabolite, ceramide, is highly conserved among eukaryotes and has been implicated in membrane organization, stress responses, and antibacterial activity [12–20]. Yet whether sphingolipid metabolism functions in vivo as an autonomous defense mechanism remains unknown. Given the emerging role of amino acids as regulators of both metabolism and immunity [21–24], we asked whether amino-acid signaling can activate sphingolipid metabolism and thereby confer resistance to infection.

The rapid and efficient adaptability of *Plutella xylostella* to diverse environmental stresses provides an ideal model system for dissecting the activation and functional characteristics of independent metabolic defense pathways [25–27]. We found that arginine activates the sphingolipid metabolic pathway in *P. xylostella*, leading to ceramide accumulation and consequent direct inhibition of pathogens. Moreover, gut bacteria can directly supply arginine to the host. This study reveals for the first time that the sphingolipid pathway can serve as a metabolic defense axis independent of classical immune pathways, such as the Toll and IMD pathways, and participate in gut defense regulation across multiple insect species. This study systematically defines an arginine–sphingolipid– ceramide defense axis, providing critical evidence for the evolutionary origins of eukaryotic defense networks and expanding new theoretical avenues for broad-spectrum anti-infection strategies that target metabolic pathways.

## Results

### Arginine mediates Bacillus thuringiensis resistance in P. xylostella

To identify amino acids capable of enhancing host defense, we first evaluated the effects of 10 amino acids (isoleucine, leucine, lysine, methionine, phenylalanine, threonine, tryptophan, valine, arginine and histidine) on the growth and development of *P. xylostella*. Phenylalanine, lysine, and leucine prolonged larval development and reduced both pupation rates and pupal weight, indicating adverse effects on host fitness (Fig 1A-1E). These amino acids were therefore excluded from subsequent infection assays. Among the remaining seven amino acids, only arginine (Arg) enhanced resistance to *Bacillus thuringiensis* (Bt) and suppressed bacterial proliferation in *vivo* (Figs 1F, S1A, and S1B). In contrast, Arg did not inhibit Bt growth in *vitro* (Fig 2A), suggesting that its protective effect is mediated through activation of host defense mechanisms rather than direct antibacterial activity.

**Fig 1.**
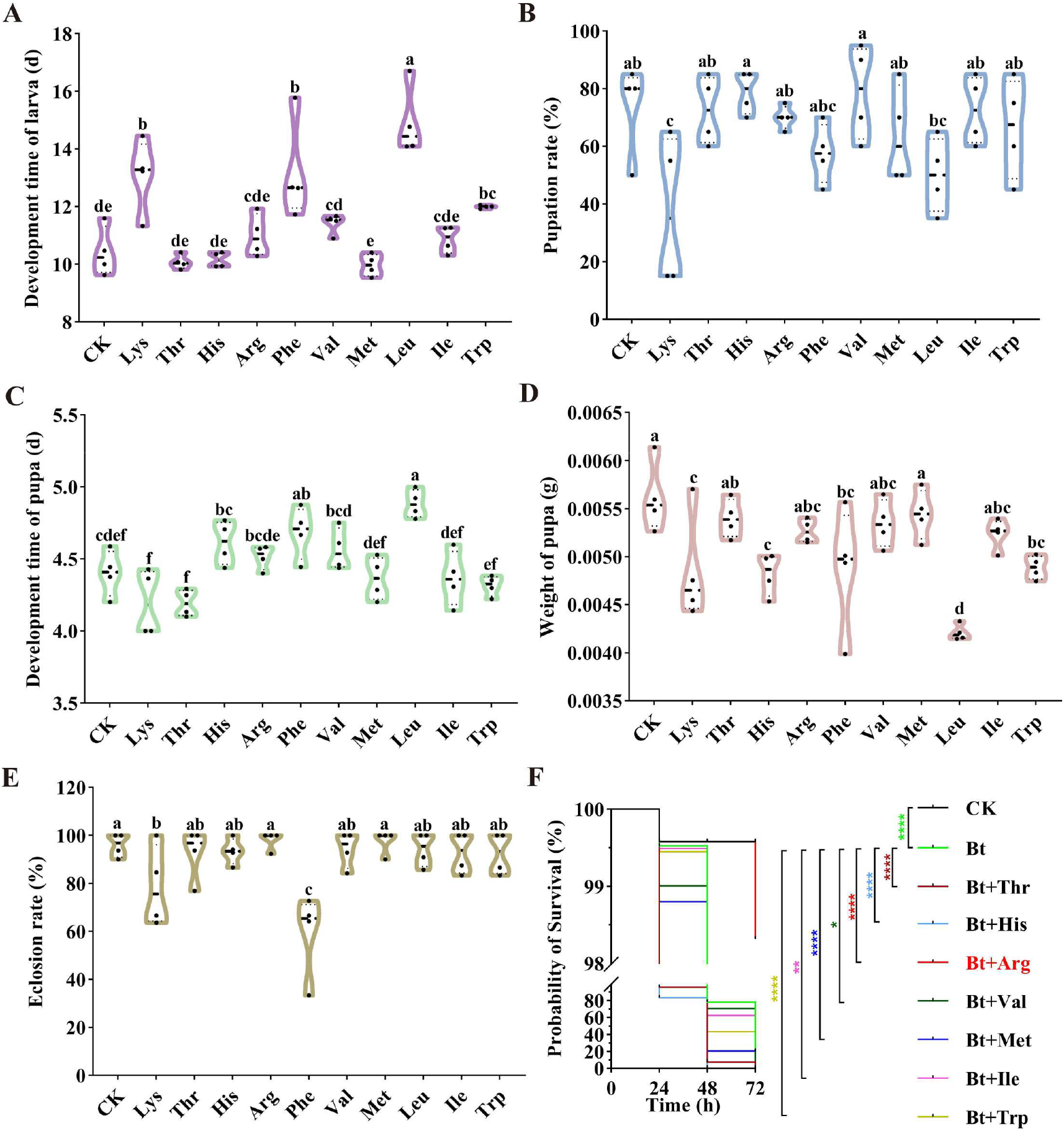
Arginine mediates Bt resistance in *P. xylostella.* (**A**) Phenylalanine, lysine, leucine, and tryptophan prolonged larval duration; n = 4 biological replicates. (**B**) Lysine reduced pupation rate; n = 4 biological replicates. (**C**) Phenylalanine and leucine prolonged pupal duration; n = 4 biological replicates. (**D**) Phenylalanine, lysine, leucine, histidine, and tryptophan reduced pupal weight; n = 4 biological replicates. (**E**) Phenylalanine and lysine reduced eclosion rate; n = 4 biological replicates. (**F**) Arginine mediated defense against Bt in *P. xylostella*; n = 4 biological replicates. Abbreviations of the full names are used in the x-axis, with the details provided in the Materials and Methods section. For (**A**) to (**E**), one-way ANOVA was performed, and different lowercase letters indicate significant differences (*P* < 0.05). For (**F**), the Mantel-Cox test was used; asterisks denote *P* <0.05(*), <0.01(**), <0.0001(****). *P* < 0.05 is considered statistically significant.

**Fig 2.**
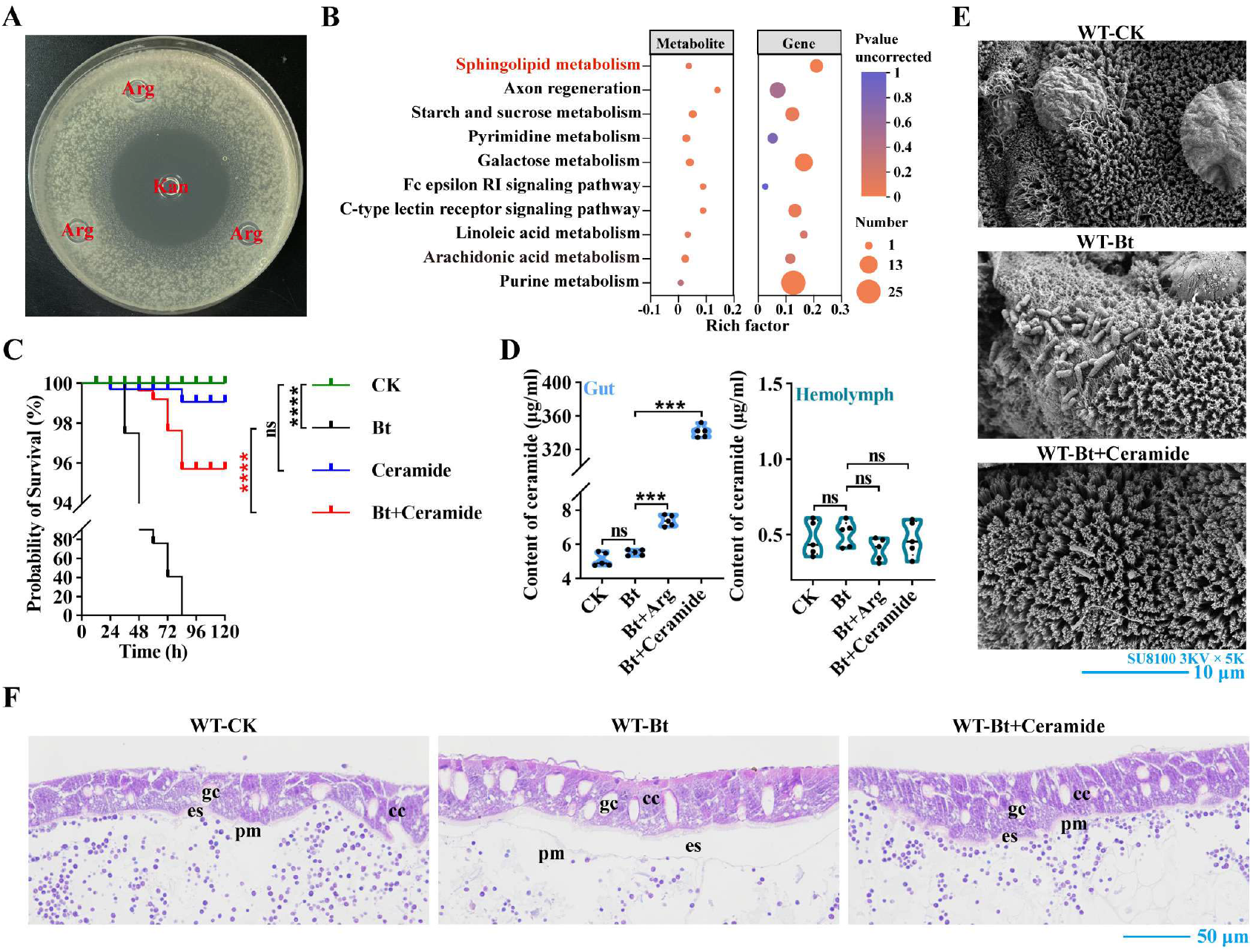
The effector ceramide in the sphingolipid metabolic pathway protects the gut of *P. xylostella.* (**A**) Arginine does not inhibit Bt growth in *vitro*. (**B**) KEGG pathway enrichment analysis of differentially expressed genes in larvae treated with arginine under Bt stress. (**C**) Ceramide mediates defense against Bt in *P. xylostella*; n = 3 biological replicates. (**D**) Arginine upregulates ceramide levels in the gut of *P. xylostella*; n = 5 biological replicates. (**E**) Ceramide protects gut microvilli from Bt damage in *P. xylostella*. (**F**) Ceramide protects the midgut peritrophic membrane from Bt damage in *P. xylostella*. WT, wild type *P. xylostella*. Abbreviations of the full names are used, with the details provided in the Materials and Methods section. gc, goblet cells; cc, columnar cells; pm, peritrophic membrane; es, space between epithelial cells and the peritrophic membrane; Kna, kanamycin. For (**C**), the Mantel-Cox test was used; asterisks indicate *P* <0.05(*), <0.0001(****), “ns” denotes no significant difference. For (**D**), the independent samples t test was used; asterisks indicate *P* <0.001(***), “ns” denotes no significant difference. *P* < 0.05 is considered statistically significant.

### Arginine induces gut ceramide synthesis to protect the gut in *P. xylostella*

To elucidate the molecular mechanism by which Arg mediates Bt resistance in *P. xylostella*, a combined transcriptomic and metabolomic analysis was performed on larvae fed with Arg under Bt infection stress. The results showed that the sphingolipid metabolic pathway was enriched (Fig 2B). Ceramide was identified as the core metabolite of this pathway. Bioassays further verified that ceramide enhanced the defense capability of *P. xylostella* against Bt (Fig 2C). Further investigation demonstrated that under Bt stress, Arg feeding significantly increased gut ceramide content (Fig 2D). Microscopic observations of the gut revealed that ceramide protected the gut microvilli and peritrophic membrane from damage caused by Bt (Fig 2E and 2F). These findings indicate that Arg induces gut ceramide synthesis in *P. xylostella*, thereby protecting the gut from Bt damage.

### Arginine mediates *Bacillus thuringiensis* resistance in *P. xylostella* via activation of the sphingolipid metabolic pathway

Under Bt stress, Arg upregulated the expression of genes involved in ceramide synthesis within the sphingolipid metabolic pathway in *P. xylostella*, the genes encoding serine palmitoyltransferase (SPT) (*Px016398*), 3-ketodihydrosphingosine reductase (KDSR) (*Px010122*), ceramide synthase (CERS) (*Px008942*), and dihydroceramide desaturase (DES) (*Px008081*) in the *de novo* synthesis pathway, as well as the genes encoding sphingomyelin phosphodiesterase 1 (SMPD1) (*Px004605* and *Px003742*) in the sphingomyelin hydrolysis pathway (S2A and S2B Fig). These results suggested that Arg activated both *de novo* synthesis and sphingomyelin hydrolysis pathways to promote ceramide production. As the key rate limiting enzyme in the *de novo* ceramide synthesis pathway, SPT and its specific inhibitor myriocin were used. Myriocin not only significantly weakened the ability of Arg to mediate Bt resistance in *P. xylostella*, but also significantly attenuated the induction of gut ceramide synthesis by Arg (S2C and S2D Fig). These findings indicated that under Bt stress, Arg activated *de novo* ceramide synthesis to mediate Bt resistance in the host.

Given that *PxDES* (*Px008081*) was a key upregulated gene at the terminal step of the *de novo* synthesis pathway, and *PxSMPD1* (*Px003742*) was a gene with significantly upregulated expression in the sphingomyelin hydrolysis pathway (S2A and S2B Fig), CRISPR/Cas9 technology was used to knock out each of these two genes, in order to further clarify the function of ceramide synthesis pathways in Bt defense of *P. xylostella*. Compared with the wild type (WT), the mutant Δ*PxDES* obtained by knocking out *PxDES* showed significantly reduced gut ceramide content after Arg feeding (Fig 3A and 3B). Bioassays revealed that under Bt stress, the survival rate of Δ*PxDES* fed with Arg was significantly lower than that of WT (Fig 3C). After ceramide supplementation, the survival rate of the mutant was not significantly different from that of WT, and the gut microvilli and peritrophic membrane remained intact without damage from Bt (Fig 3C-3E). These results indicated that deletion of *PxDES* prevented Arg from activating *de novo* ceramide synthesis pathway, thereby abolishing the Bt defense function in *P. xylostella*. The mutant Δ*PxSMPD1* obtained by knocking out *PxSMPD1* exhibited significantly less gut ceramide content after Arg feeding under Bt stress than WT (S3A and S3B Fig). The survival rate of Δ*PxSMPD1* was significantly decreased, but after ceramide supplementation, the survival rates of the mutant and WT did not significantly differ (S3C and S3D Fig). These findings demonstrated that deletion of *PxSMPD1* significantly inhibited the activation of the sphingomyelin hydrolysis pathway by Arg, thereby attenuating the ability of Arg to mediate Bt resistance in *P. xylostella*. Collectively, these results confirmed that Arg activated sphingolipid metabolism to promote ceramide synthesis, thereby mediating the defense of *P. xylostella* against Bt.

**Fig 3.**
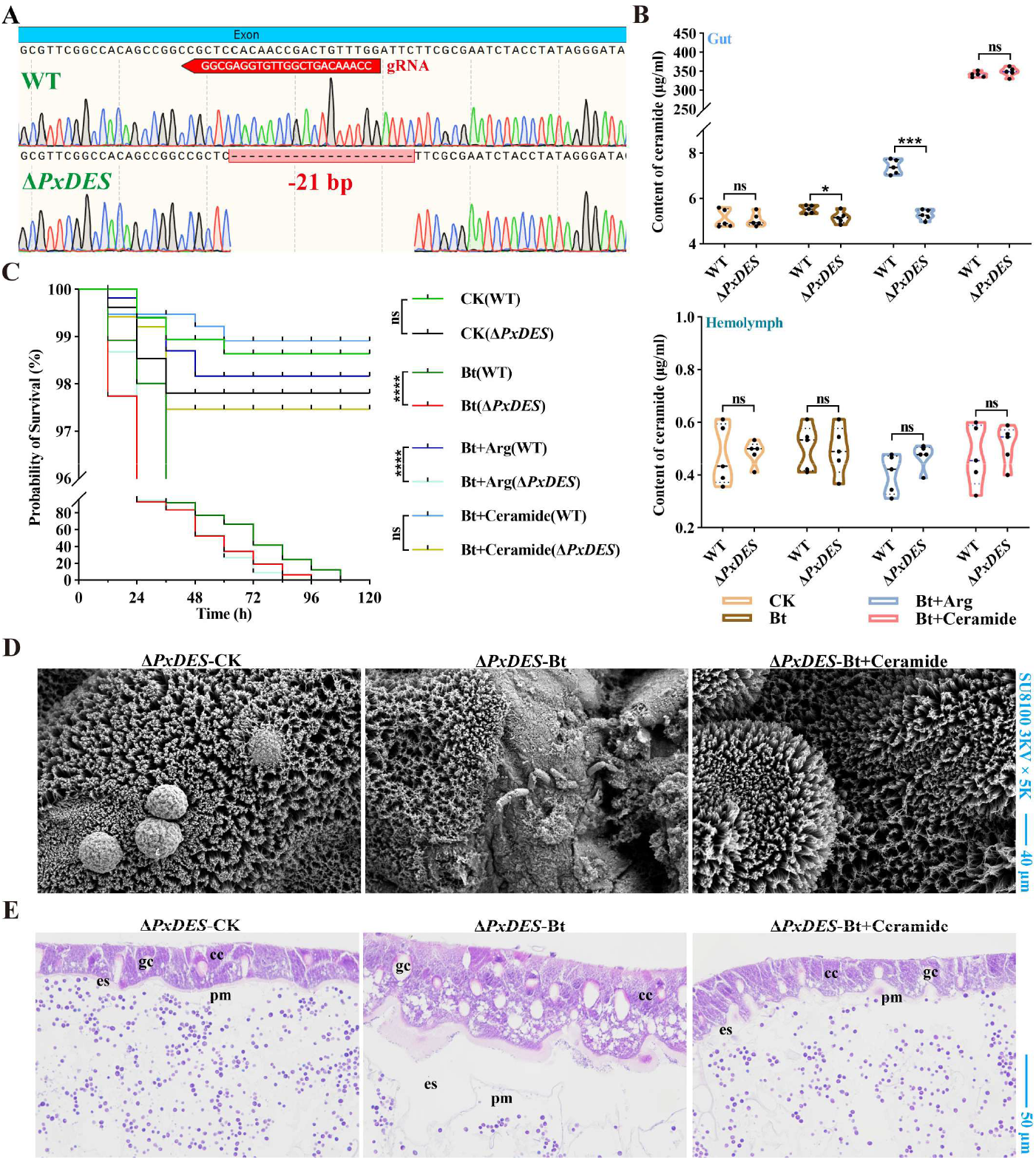
The *de novo* ceramide synthesis pathway gene *PxDES* mediates defense against Bt in *P. xylostella.* (**A**) Generation of the mutant Δ*PxDES* in *P. xylostella*; the mutant carries a deletion of 21 bp. (**B**) Deletion of *PxDES* significantly attenuates the arginine induced increase in gut ceramide levels; n = 5 biological replicates. (**C**) Deletion of *PxDES* abolishes the protective effect of arginine against Bt in *P. xylostella*; n = 3 biological replicates. (**D**) Arginine fails to protect gut microvilli from Bt damage in the Δ*PxDES* mutant. (**E**) Ceramide protects the midgut peritrophic membrane from Bt damage in the Δ*PxDES* mutant. WT, wild type; Δ*PxDES*, the mutant carrying a deletion of *PxDES*. Abbreviations of the full names are used, with the details provided in the Materials and Methods section. gc, goblet cells; cc, columnar cells; pm, peritrophic membrane; es, space between epithelial cells and the peritrophic membrane. For (**B**), the independent samples t test was used; asterisks indicate *P* <0.05(*), <0.001(***), “ns” denotes no significant difference. For (**C**), the Mantel-Cox test was used; asterisks indicate *P* < 0.0001 (****), “ns” denotes no significant difference. *P* < 0.05 is considered statistically significant.

### Ceramide exerts antibacterial effects as a direct inhibitory factor

To determine whether ceramide functions as a second messenger in mediating insect resistance against pathogenic bacteria, transcriptomic analysis was performed on *P. xylostella* fed with ceramide under Bt stress. The results revealed that the classical defense pathways, namely the Toll and IMD signaling pathways, were not significantly enriched (P adjusted = 0.100), whereas the peroxisome pathway was significantly enriched (P adjusted = 0.002) (Fig 4A). Peroxisomes are core functional units for the production and scavenging of reactive oxygen species (ROS). However, ROS detection showed that under Bt stress, gut ROS levels in *P. xylostella* fed with either Arg or ceramide did not increase significantly (Fig 4B and 4C). Neither Arg nor ceramide induced the upregulation of antimicrobial peptide genes, including *Cecropin2*, *Gloverin2*, *Lyszomes2*, and *Moricin3* (Fig 4D-4G). These results indicated that neither Arg nor ceramide could induce ROS or antimicrobial peptide mediated Bt resistance in *P. xylostella*.

**Fig 4.**
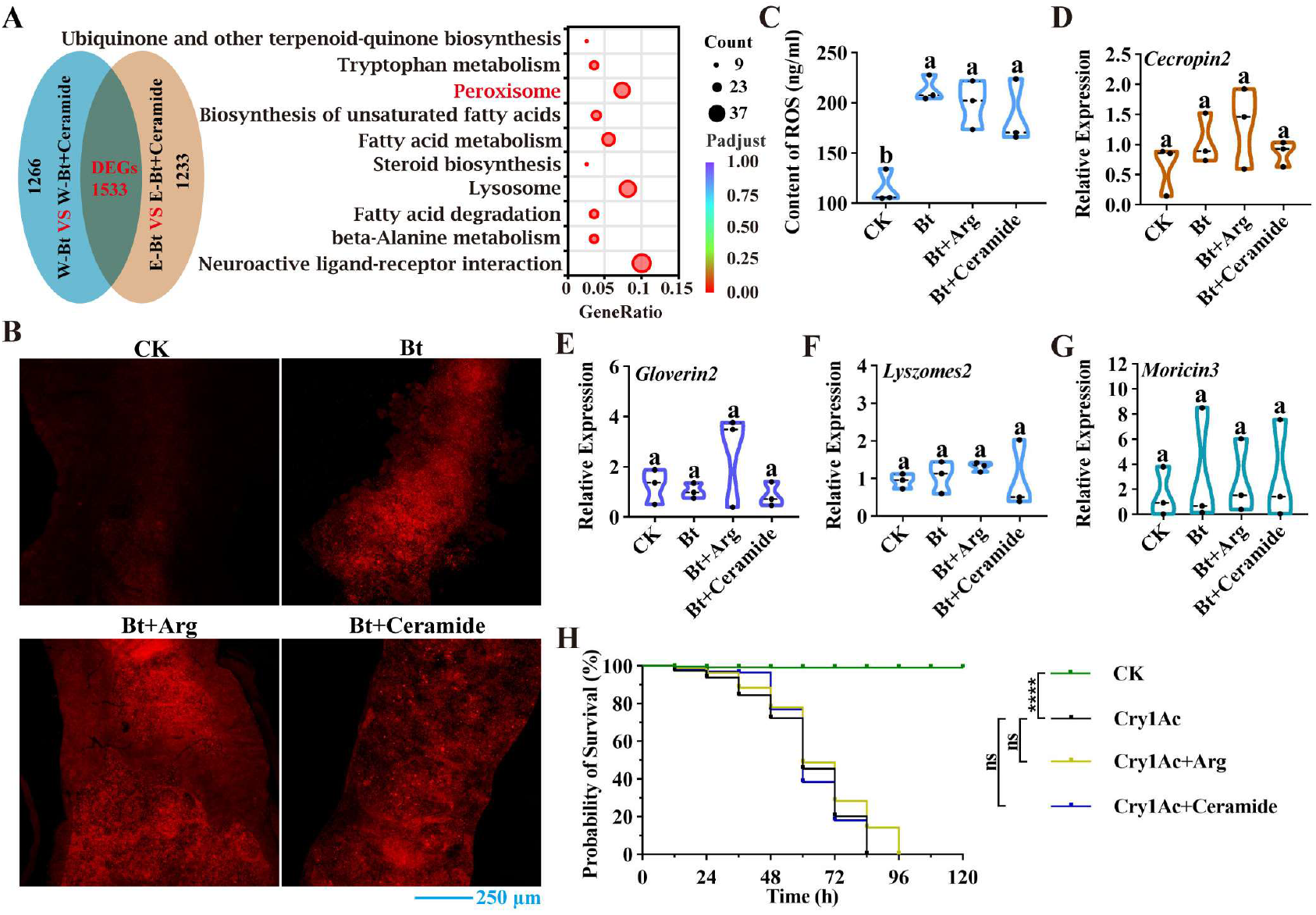
Ceramide does not induce reactive oxygen species or antimicrobial peptide mediates host defense against Bt. (**A**) Transcriptomic analysis of larvae treated with ceramide. Venn analysis of differentially expressed genes (DEGs) (left panel), joint KEGG enrichment analysis of DEGs (right panel). (**B**) Ceramide does not upregulate reactive oxygen species levels in gut (DHE staining). (**C**) Ceramide does not induce reactive oxygen species synthesis in gut (quantitative detection); n = 3 biological replicates. (**D**) Ceramide does not upregulate the expression of the antimicrobial peptide gene *Cecropin2*; n = 3 biological replicates. (**E**) Ceramide does not upregulate the expression of the antimicrobial peptide gene *Gloverin2*; n = 3 biological replicates. (**F**) Ceramide does not upregulate the expression of the antimicrobial peptide gene *Lyszomes2*; n = 3 biological replicates. (**G**) Ceramide does not upregulate the expression of the antimicrobial peptide gene *Moricin3*; n = 3 biological replicates. (**H**) Arginine and ceramide do not assist *P. xylostella* in defending against the Bt toxin Cry1Ac; n = 4 biological replicates. WT, wild type; Δ*PxDES*, the mutant carrying a deletion of *PxDES*. Abbreviations of the full names are used in the x-axis, with the details provided in the Materials and Methods section. For (**C**) to (**G**), one-way ANOVA was performed, and different lowercase letters indicate significant differences (*P* < 0.05). For (**H**), the Mantel-Cox test was used; asterisks indicate *P* < 0.0001 (****), “ns” denotes no significant difference. *P* < 0.05 is considered statistically significant.

Bt primarily exerts its insecticidal effect by producing specific insecticidal toxins and proteins that target the midgut epithelial cells of insects. Nevertheless, experiments demonstrated that neither Arg nor ceramide could mediate resistance of *P. xylostella* against the Bt toxin Cry1Ac (Fig 4H), suggesting that ceramide may exert its function through direct inhibition of Bt cells. In *vitro* experiments confirmed that ceramide directly inhibited Bt growth, with a minimum inhibitory concentration (MIC) of 2.5 mg/mL (Fig 5A). With increasing concentrations of ceramide, Bt not only failed to form parasporal crystals, but the disruption of vegetative cells also increased in a concentration dependent manner (Fig 5B). Transmission electron microscopy observations showed that ceramide disrupted the cell wall structure of Bt, leading to leakage of cytoplasmic contents (Fig 5C). Broad-spectrum antibacterial validation demonstrated that ceramide exerted inhibitory effects against a range of bacteria, including *Lactobacillus carnis*, *Staphylococcus aureus*, and *Serratia marcescens*, with more pronounced inhibition against Gram-positive bacteria (S4A-S4C Fig). The antibacterial mechanism of ceramide appeared to be species specific. In *S. marcescens* (Gram-negative), it inhibited vegetative cell division and caused cell elongation, whereas in *S. aureus* (Gram-positive), it directly caused disruption of vegetative cells (S4D and S4E Fig). These observations suggested that the antibacterial mode of ceramide may differ among bacterial species. Further analysis via 16S rRNA amplicon sequencing of gut bacteria from *P. xylostella* revealed that ceramide alleviated the disturbance of gut bacterial richness and diversity caused by Bt, without affecting overall gut microbial diversity (Fig 5D). This finding indicated that ceramide exhibited relatively high specificity in inhibiting pathogenic bacteria. Collectively, these results confirmed that ceramide functions as a direct inhibitory factor to mediate resistance against pathogenic bacteria in *P. xylostella*.

**Fig 5.**
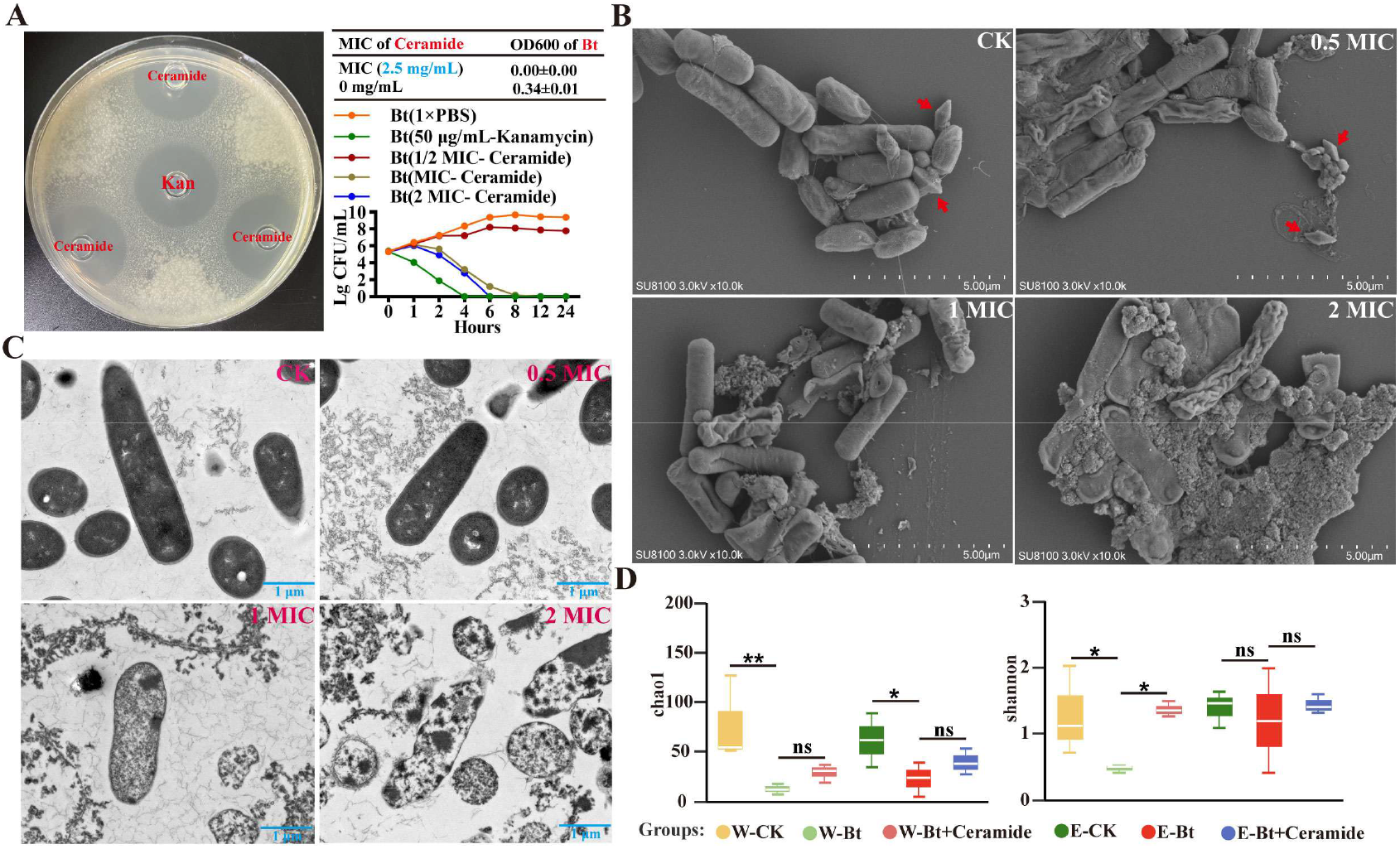
Ceramide disrupts the Bt cell wall and consequently causes cytoplasmic efflux. (**A**) In *vitro* inhibition of Bt by ceramide. The plate assay shows the results of the Oxford cup inhibition zone method [51]; the three-line table presents the minimum inhibitory concentration (MIC) of ceramide against Bt; the curve graph displays the inhibition kinetics of Bt at different MICs of ceramide; n = 3 technical replicates. (**B**) Ceramide directly disrupts the vegetative cell structure of Bt. Red arrows indicate the parasporal crystals of Bt. (**C**) Ceramide damages the Bt cell wall and consequently causes cytoplasmic dissolution. (**D**) Ceramide alleviates the disruption of gut bacterial homeostasis in *P. xylostella* caused by Bt. WT, wild type; Δ*PxDES*, the mutant carrying a deletion of *PxDES*. Abbreviations of the full names are used, with the details provided in the Materials and Methods section. Kna, kanamycin; CK, control without ceramide treatment; 0.5 MIC, 1 MIC, and 2 MIC represent different concentrations of ceramide relative to the MIC. For (**D**), the nonparametric Kruskal-Wallis rank sum test was used, asterisks indicate *P* <0.05(*), <0.01(**), “ns” denotes no significant difference. *P* < 0.05 is considered statistically significant.

### Ceramide antipathogenic effect shows potential universality across insect species

To determine whether ceramide exhibits broad-spectrum activity against different types of agents, challenges with *Serratia marcescens* (a Gram-negative pathogenic bacterium, Sm), *Beauveria bassiana* (a pathogenic fungus), and the chemical insecticides cyfluthrin and dipterex were tested in *P. xylostella*. The results showed that ceramide could not mediate resistance of *P. xylostella* against cyfluthrin, dipterex, or *B. bassiana* (S5A-S5C Fig). However, ceramide significantly enhanced resistance against *S. marcescens* (S5D Fig), indicating that ceramide specifically mediates resistance against pathogenic bacteria in *P. xylostella*.

The key enzymes for ceramide synthesis in the sphingolipid metabolic pathway, including SPT, KDSR, CERS, DES, SMPD, and GBA, were found not only in *P. xylostella* but also in the model insect *Drosophila melanogaster*, as revealed by Basic Local Alignment Search Tool of National Center for Biotechnology Information (NCBI BLAST) searches. Notably, these enzymes are also present in other eukaryotes such as *Mus musculus*, *Danio rerio*, *Homo sapiens*, *Sus scrofa* and *Gallus gallus* (S6A Fig), indicating their evolutionary conservation across eukaryotes. To investigate whether the antipathogenic effect of ceramide is broadly applicable across insect species, *D. melanogaster* and *Tuta absoluta* were selected as test subjects. Bioassay results showed that *D. melanogaster* was not significantly sensitive to Bt, whereas ceramide assisted *T. absoluta* in defending against Bt (S6B and S6C Fig). Under Sm challenge, ceramide enhanced resistance against Sm in both *T. absoluta* and *D. melanogaster* (S6D and S6E Fig). These findings indicate that the antipathogenic effect of ceramide exhibits potential universality across insect species.

### Arginine produced by gut bacterium *Raoultella ornithinolytica* induces ceramide synthesis to mediate *Bacillus thuringiensis* resistance in *P. xylostella*

Arg is a conditionally essential amino acid that must be acquired exogenously by *P. xylostella*. Previous studies have shown that gut microbiota can supply amino acids to the host. In the present study, it was found that the gut bacterium *Raoultella ornithinolytica* (OV6) of *P. xylostella* efficiently synthesizes Arg (Fig 6A), and under Bt stress, Arg content in the gut significantly increased after feeding with OV6 (Fig 6B). Further bioassay validation demonstrated that under Bt stress, WT *P. xylostella* fed with OV6 exhibited significantly increased survival rates, whereas the Δ*PxDES* mutant fed with OV6 showed significantly lower survival rates than the wild type (Fig 6C). OV6 did not exhibit antibacterial activity against Bt in *vitro* (Fig 6D), suggesting that gut bacterium OV6 may mediate host defense against Bt through Arg synthesis.

**Fig 6.**
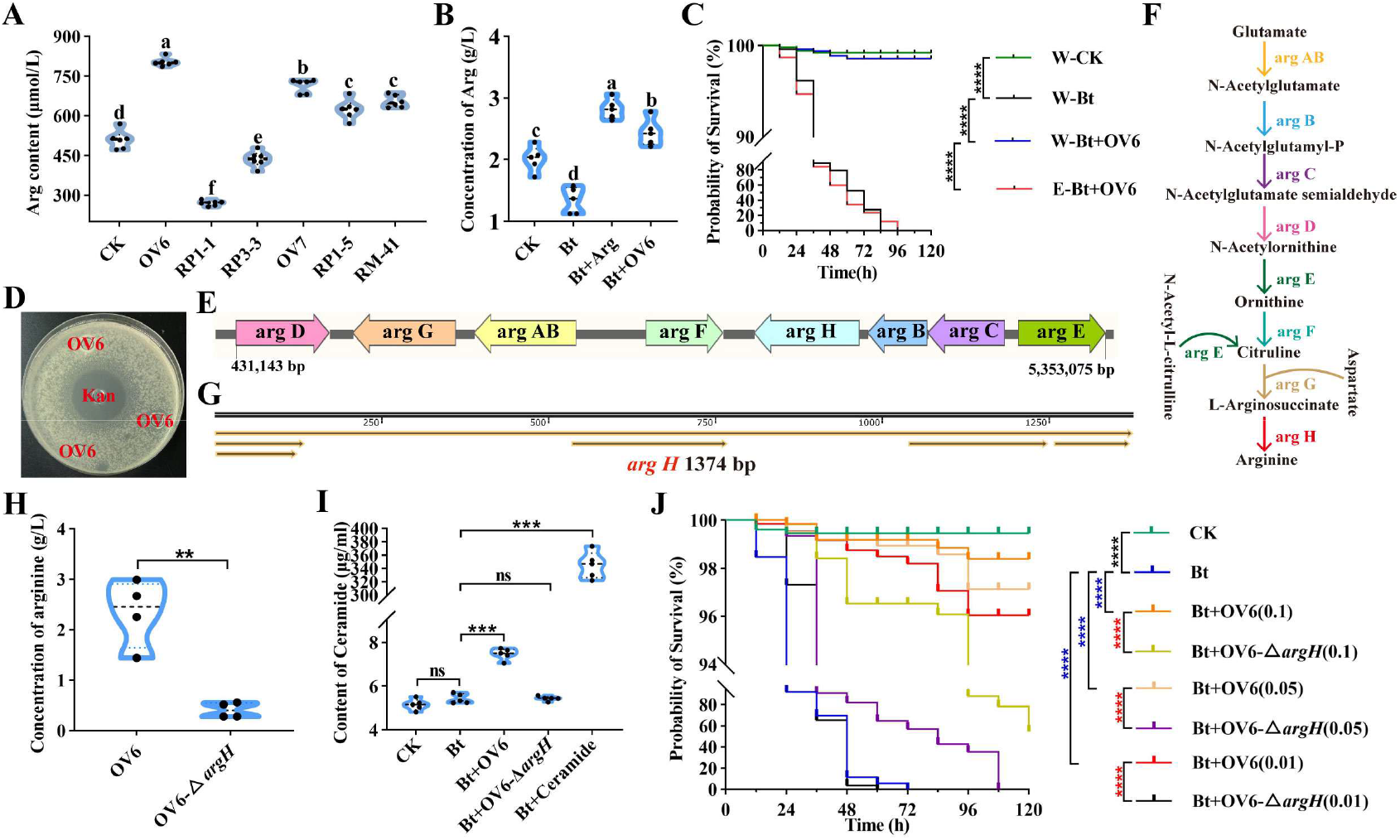
Arginine from gut bacterium OV6 induces host ceramide synthesis to confer Bt resistance in *P. xylostella.* (**A**) Gut bacterium OV6 efficiently synthesizes arginine; n = 6 technical replicates. (**B**) Gut bacterium OV6 upregulates arginine content in the gut of *P. xylostella*; n = 5 biological replicates. (**C**) Gut bacterium OV6 mediates host defense against Bt in wild type *P. xylostella*; n = 3 biological replicates. (**D**) Gut bacterium OV6 does not inhibit Bt growth in *vitro*. (**E**) Schematic diagram of the complete gene cluster for arginine synthesis in gut bacterium OV6. (**F**) Pathway map of arginine synthesis in gut bacterium OV6. (**G**) The key gene *argH* in the arginine synthesis pathway of gut bacterium OV6 is 1374 bp in size. (**H**) Arginine production is significantly reduced in the OV6-Δ*argH* mutant strain; n = 4 technical replicates. (**I**) The ability of the OV6-Δ*argH* mutant strain to induce gut ceramide synthesis in *P. xylostella* is significantly reduced; n = 5 biological replicates. (**J**) The OV6-Δ*argH* mutant strain loses its ability to mediate host defense against Bt in *P. xylostella*; n = 4 biological replicates. WT, wild type; Δ*PxDES*, the mutant carrying a deletion of *PxDES*. Abbreviations of the full names are used, with the details provided in the Materials and Methods section. Kna, kanamycin. For (**A**) and (**B**), one-way ANOVA was performed, and different lowercase letters indicate significant differences (*P* < 0.05). For (**C**) and (**J**), the Mantel-Cox test was used; asterisks indicate *P* < 0.0001(****), “ns” denotes no significant difference. For (**H**) and (**I**), the independent samples t test was used, asterisks indicate *P* < 0.01(**), < 0.001(***), “ns” denotes no significant difference. *P* < 0.05 is considered statistically significant.

Genomic analysis confirmed that OV6 possesses a complete Arg biosynthesis gene cluster (Fig 6E), with *argH* located at the terminal step of the Arg synthesis pathway (Fig 6F and 6G). To determine whether Arg derived from OV6 synthesis mediates host defense against Bt, the key gene *argH* was knocked out to obtain the mutant strain OV6-Δ*argH* (S7 Fig). Compared with the wild type OV6, the Arg production of OV6-Δ*argH* was significantly reduced (Fig 6H). Functional validation showed that in *P. xylostella* fed with wild type OV6, gut ceramide content was significantly elevated, whereas no significant change was observed in the gut ceramide content of larvae fed with OV6-Δ*argH* (Fig 6I). OV6-Δ*argH* at different concentrations failed to mediate host defense against Bt (Fig 6J). These results indicate that gut bacterium OV6 mediates defense against Bt in *P. xylostella* by synthesizing Arg, which in turn induces ceramide synthesis.

## Discussion

In this study, we demonstrate that gut bacterium-derived arginine functions as a key metabolic signal that activates the conserved sphingolipid pathway, promotes ceramide accumulation, and enhances host resistance to infection. Our findings establish the sphingolipid pathway as an autonomous metabolic defense module that operates independently of canonical Toll and IMD immune signaling. Furthermore, we show that ceramide acts as a direct antibacterial effector with preferential activity against Gram-positive bacteria. Together, these results reveal a previously unrecognized mechanism by which bacterium-derived metabolites engage host metabolic pathways to mediate innate immune defense. More broadly, our study supports the emerging view that metabolism is not merely a source of energy and biosynthetic intermediates for immune responses, but can also function as a direct determinant of pathogen resistance.

Sphingolipid metabolism constitutes a complex network of interconnected metabolic pathways, and its products including ceramide, sphingosine, and sphingosine 1 phosphate (S1P) play important roles in regulating cell growth, death, senescence, adhesion, migration, inflammation, angiogenesis, and intracellular trafficking [28, 29]. Notably, the immune defense effects of the sphingolipid metabolic pathway were found to be analogous to those of the classical Toll and IMD pathways, jointly forming a “dual assurance” system of gut immunity. Both possess specific antibacterial activities: the Toll pathway preferentially targets Gram-positive bacteria, while the IMD pathway primarily responds to Gram-negative bacteria [30]. The core effector of sphingolipid metabolism, ceramide, mainly inhibits Gram-positive bacteria, and this targeted antibacterial feature echoes the pathway specificity of Toll and IMD. Both also maintain gut microbial homeostasis: the Toll and IMD pathways specifically eliminate foreign bacteria and pathogens by activating defense responses such as antimicrobial peptides and reactive oxygen species [31, 32]. This study demonstrated that ceramide, while exerting its antibacterial effects, effectively alleviates the disturbance of gut bacterial diversity in *P. xylostella* under Bt stress. Given that ceramide acts primarily on Gram-positive bacteria, and that proteobacteria, all of which are Gram-negative, constitute the dominant phylum in the guts of various insects (e.g., *Cnaphalocrocis medinalis*, *Bactrocera minax*, and *P. xylostella*) [33–35], this finding provides a reasonable explanation for the specific regulatory mode of ceramide on the gut microbiota. In summary, the sphingolipid metabolic pathway, like the Toll and IMD pathways, plays important roles in both “anti-infection” and “maintenance of commensal homeostasis” in the host.

The ceramide synthesis pathway within sphingolipid metabolism is highly conserved across eukaryotes. In mammals, ceramide functions by inducing apoptosis, mediating autophagy, regulating fatty acid metabolism, and modulating immunity [15–18]. In insects, ceramide also undertakes important physiological functions, such as serving as a major component of the cuticle in *Tribolium castaneum*, participating in lipid transport and construction of the body surface barrier [36]. Notably, ceramide possesses direct antibacterial activity: ceramide and its analogues can inhibit the activity of the Gram-negative bacteria *Neisseria* sp. and *Acinetobacter baumannii* [19, 20]. Molecular docking has shown that ceramide can interact with membrane proteins and biofilm associated enzymes of *A. baumannii* through hydrogen bonding of functional groups, thereby disrupting membrane permeability [20]. The present study further complements the mechanism by which ceramide inhibits Gram-negative bacteria, revealing that it can inhibit the division process of vegetative cells of Gram-negative bacteria, resulting in an abnormally elongated phenotype. This phenomenon may arise from differences in the antibacterial mechanisms of ceramide against different Gram-negative bacteria, or from the coexistence of the above-mentioned membrane permeability disruption and proliferation inhibition mechanisms. At the same time, the specific mechanism by which ceramide targets the cell membranes of Gram-negative bacteria remains to be further verified. In addition, this study found that ceramide exerts particularly pronounced inhibitory effects on Gram positive bacteria, being able to disrupt their cell wall structure and cause leakage of cytoplasmic contents. The reason for the mechanistic specificity of ceramide against Gram-negative versus Gram-positive bacteria may be as follows: the outer membrane of Gram-negative bacteria serves as the first barrier [37]; ceramide may not directly act on the thin peptidoglycan layer inside, but may first bind to membrane proteins and biofilm associated enzymes on the outer membrane through hydrogen bonding of functional groups, disrupt membrane permeability, and then further inhibit vegetative cell division. Gram-positive bacteria lack an outer membrane barrier [37]; ceramide can directly contact and disrupt the dense peptidoglycan layer, leading to cell wall structural damage and cytoplasmic leakage. Some biological control strategies exhibit host or environmental dependence, and medical anti-infection therapies are often limited by species specific immune mechanisms [38, 39]. In contrast, intervention strategies based on the sphingolipid metabolic pathway could be applied across species. In agriculture, by regulating arginine synthesis in pest gut bacteria or directly delivering ceramide synthesis inhibitors, broad-spectrum green control technologies could be developed against multiple pests such as *P. xylostella* and *Tuta absoluta*. In the medical field, given the inhibitory effect of ceramide against human pathogens such as *Staphylococcus aureus*, modulation of gut bacterial Arg metabolism or sphingolipid synthesis could provide novel therapeutic approaches for drug resistant bacterial infections. Furthermore, artificial synthesis and application of ceramide analogues could be explored to optimize antibacterial activity and stability, developing new antibacterial agents applicable in both agriculture and medicine.

The present study reveals that the key metabolic signal Arg in the Arg, sphingolipid metabolism, pathogen resistance network originates from the gut bacterium *Raoultella ornithinolytica* (OV6) of *P. xylostella*, which is widely distributed in soil and water environments and can also form symbioses with multiple eukaryotes [40]. Previous reports on the regulation of host disease resistance by gut microbiota have largely focused on mechanisms such as siderophore production, toxin degradation, or induction of immune pathways [41, 42]. This study expands the known mechanisms by which gut bacteria regulate host disease resistance, demonstrating that OV6, as an insect gut bacterium, can activate the sphingolipid metabolic pathway through its own metabolite Arg, thereby mediating host defense against pathogens. These finding echoes related studies in *Aedes albopictus*, where the gut bacterium *Enterobacter hormaechei* B17 (Eh_B17) secretes sphingosine that directly inhibits Zika and Dengue viruses in the host [43], with the core being that the symbiont directly produces the lipid molecule D erythro sphingosine. The present study demonstrates that gut bacteria can activate host sphingolipid metabolism to synthesize ceramide, which exerts direct antibacterial effects. These two regulatory modes represent “direct supply” and “indirect activation,” respectively, and together constitute a dual mechanism by which gut bacteria regulate the lipid metabolic network.

This study elucidates the molecular mechanism by which the gut bacterium OV6 derived metabolite Arg activates the sphingolipid metabolic pathway to mediate defense against pathogens in *P. xylostella* and also expands the understanding of gut bacteria mediated host disease resistance. Nevertheless, the regulatory mechanism by which Arg modulates the sphingolipid metabolic pathway remains to be further dissected. In organisms, Arg supply is achieved through multiple transport systems that uptake extracellular Arg, including the cationic amino acid specific transport system y^+^ and several broad amino acid transport systems (y^+^L, b^0,+^ and B^0,+^) [44]. Furthermore, amino acid transporters with system y^+^ characteristics (CATs) primarily transport cationic amino acids such as arginine through facilitated diffusion and differential trans stimulation of intracellular substrates [45]. Future studies could clarify and elucidate the transport mechanism of Arg from the gut into cells through the aforementioned amino acid transport systems. Previous studies have confirmed that Arg can activate the GPRC6A/PI3K/AKT/mTOR signaling pathways [46, 47], and the mTORC1-S6K1-SRPK2 signaling pathway significantly regulates lipid biosynthesis [48]. On the basis of clarifying the Arg transport mechanism, further investigation could explore whether Arg regulates the transcriptional expression of key genes in the sphingolipid metabolic pathway through the GPRC6A/PI3K/AKT/mTOR signaling axis, thereby completing the upstream regulatory mechanism of Arg mediated sphingolipid activation. Nonetheless, this study elucidates the molecular mechanism by which Arg derived from gut bacterium OV6 activates the sphingolipid metabolic pathway to mediate defense against pathogens in *P. xylostella*, revealing that the insect mobilizes a basal metabolic pathway directly as an immune effector system, rendering small molecule metabolites as immune effector factors on par with antimicrobial peptides and reactive oxygen species. This finding will elevate the traditional understanding of insect immunity as essentially equivalent to protein based cellular immunity, adding a new metabolic dimension to the theory of multi module network immunity. Moreover, the elucidation of the insect metabolic defense module indicates that organisms had already evolved strategies to utilize basal metabolic pathways for immune defense prior to the emergence of classical immunity, which may represent a more ancient and fundamental defense mode than protein-based immunity, thereby expanding the understanding of insect innate immune defense.

## Materials and Methods

### Insect Rearing

*P. xylostella* was reared in an artificial climate chamber under conditions of 70-80 % relative humidity, temperature 25±1 ℃, and a 16 h light/8 h dark photoperiod. Larvae were fed a modified artificial diet (optimized based on the original formula [49]: wheat germ powder content reduced to 1/4 of the original formula, and the contents of methylparaben, sorbic acid, multivitamins, vitamin C, linseed oil, and linoleic acid were all reduced to 1/10 of the original formula); adults were fed 10 % honey aqueous solution. The purpose of the modification is to minimize the interference of amino acids in the feed on this study.

*D. melanogaster* was reared in an artificial climate chamber under conditions of 55-65 % relative humidity, temperature 25±1 ℃, and a 12 h light/12 h dark photoperiod. Standard cornmeal-sugar-agar medium (1 L): 105 g glucose, 7.5 g agar, 26 g yeast, 50 g cornmeal, 0.25 g sodium benzoate, 8.5 mL 95 % ethanol, and 1.9 mL 99 % propionic acid.

*T. absoluta* was reared in an artificial climate chamber under conditions of 70-80 % relative humidity, temperature 25±1 ℃, and a 16 h light/8 h dark photoperiod. Tomato seeds (Provence) were cultivated into tomato seedlings under conditions of 26 ℃, light intensity of 3000-5000 lx, and a 12 h light/12 h dark photoperiod for *T. absoluta* feeding.

### Bacterial Culture

Strains preserved at –80 ℃ were streaked onto Lysogeny Broth (LB) solid medium and cultured for 12 h. Single colonies were picked and inoculated into LB liquid medium, then cultured in a constant temperature shaker at 180 rpm for 12 h for activation: the culture temperature for *Raoultella ornithinolytica* (OV6), *Raoultella ornithinolytica*-Δ*ArgH* (OV6-Δ*ArgH*), and *Staphylococcus aureus* (Sa) was 37 ℃, while that for *Bacillus thuringiensis* 8010 (Bt) and *Serratia marcescens* (Sm) was 30 ℃. After activation, the bacterial solution was centrifuged at 6000 rpm for 10 min at 4 ℃ to collect the bacterial cells, which were resuspended in 100 μL 1×PBS, washed by centrifugation 3 times repeatedly, and finally resuspended in 200 μL 1×PBS to determine the OD_600_ value for later use.

### Culture of *Beauveria bassiana* and preparation of spore suspension

*Beauveria bassiana* (Bb202) blocks preserved at –80 ℃ were inoculated onto Potato Dextrose Agar (PDA) solid medium and activated by culturing at 25 ℃ for 5 d; activated blocks were transferred to fresh PDA solid medium and cultured at 25 ℃ for 3 d. Conidia were collected by rinsing with 5 mL 0.05 % Tween-80 solution in an ultra-clean workbench to obtain the spore suspension. The suspension was filtered through 4 layers of sterile lens paper and transferred to a sterile 15 mL centrifuge tube. The concentration was adjusted to the required experimental concentration by counting with a hemocytometer.

### Treatment of artificial diet

Each treatment group in the experiment corresponded to one type of *P. xylostella* diet: CK treatment group (normal diet); single amino acid treatment group (Lysine (Lys), Threonine (Thr), Histidine (His), Arginine (Arg), Phenylalanine (Phe), Valine (Val), Methionine (Met), Leucine (Leu), Isoleucine (Ile), Tryptophan (Trp); concentration all 5 mg/mL); single microorganism treatment group (OD_600_ of OV6, Bt, and Sm were 0.1, 0.015, and 0.03 respectively; concentration of Bb202 spores was 5.08×10⁷ spores/mL); single toxin/insecticide treatment group (concentrations of Cry1Ac, Cyfluthrin, and Dipterex were 20 μg/mL, 1.52 mM, and 687.33 μM respectively); single other substance treatment group (concentrations of ceramide and Myriocin (M), were 5 mg/mL and 50 μM respectively); combined treatment group (Bt+single amino acid, Bt+OV6, Bt+OV6-Δ*argH*, Bt+Ceramide, Bt+M, Bt+Arg+M, Cry1Ac+Arg, Cry1Ac+Ceramide, Bb202+Ceramide, Sm+Ceramide, Cyfluthrin+Ceramide, Dipterex+Ceramide; concentrations were the same as those of the above single substances). In addition, in the bioassay study of the “OV6 synthesizes Arg to mediate Bt resistance in *P. xylostella*” result section, OV6 and OV6-Δ*argH* were both set with OD_600_ concentration gradients: 0.1, 0.05, 0.01. All diets were freshly prepared and replaced every 24 h; in the *D. melanogaster* bioassay study, the OD_600_ of Bt and Sm were 0.015 and 1 respectively, and the ceramide concentration was 5 mg/mL; in the *T. absoluta* bioassay study, the OD_600_ of Bt and Sm used to soak tomato seedlings were both 0.6, and the ceramide concentration was 5 mg/mL.

### Fitness determination of *P. xylostella*

Determination of fitness of wild-type (WT) *P. xylostella* under amino acid treatment: the normal diet, diets containing individual amino acids, with 4 replicates per group and 20 newly hatched larvae per replicate. The larval stage, pupal stage, pupal weight, pupation rate, and eclosion rate were recorded.

### Resistance-related bioassays

Each diet or soaking solution corresponded to 1 treatment group, with 3 or more biological replicates set for each treatment group (specific descriptions of biological replicates in different bioassay studies are shown in the figure legends of this study), and 20 larvae per replicate. The number of survivors was recorded.

### Detection of Arginine production by *P. xylostella* gut bacteria

#### (1) Preparation and derivatization of Arg standard solution

The 2.5 mmol/L Arg standard stock solution was diluted with ultrapure water to a series of standard solutions of 62.5 µmol/L, 125 µmol/L, 250 µmol/L, 500 µmol/L, and 1000 µmol/L. 70 µL borate buffer and 10 µL standard solution were sequentially added to a 1.5 mL centrifuge tube, mixed, then 20 µL AccQ•Tag reagent was added, vortexed for 20 s, incubated in a water bath at 55 ℃ for 10 min, cooled for 3 min, vortexed for 20 s, and transferred to a sample vial for analysis.

#### (2) Preparation and derivatization of sample solution

After activating the 6 bacterial strains OV6, *Carnobacterium gallinarum* (OV7), *Kocuria palustris* (RP1-1), *Enterococcus* sp. (RP1-5), *Microbacterium arborescens* (RP3-3), and *Brachybacterium* sp. (RM-41), 1 mL bacterial solution was inoculated into 100 mL LB liquid medium, cultured at 37 ℃ and 180 rpm for 72 h. 1.5 mL bacterial solution was centrifuged at 6000 rpm for 10 min at 4 ℃, 500 µL supernatant was taken into a 1.5 mL centrifuge tube, 500 µL acetonitrile was added, vortexed for 30 s, stood at 4 ℃ for 10 min, centrifuged at 13000 rpm for 20 min, stood for 20 min, 400 µL supernatant was filtered through a 0.22 µm filter membrane, and then treated according to the arginine standard derivatization method, with 6 replicates per strain.

#### (3) Chromatographic detection and data processing

Chromatographic conditions were referenced and optimized from the paper [50]: column temperature 49 ℃, elution gradient see S1Table. UNIFI software was used for automatic identification and integration, and manual verification was performed. A standard curve was established by linear regression of the peak area of the standard against concentration, and the Arg concentration was calculated by substituting the sample peak area.

### In *Vivo* inhibitory effects of Arg on Bt

We prepare two types of diets: one containing only Bt and the other containing Bt plus Arg. The OD_600_ of Bt is 0.015, and the Arg concentration is 5 mg/mL. Each diet is treated as one experimental group, with three replicates per group and 15 larvae per replicate. Third-instar late-stage larvae of *P. xylostella* reared on normal diet were selected, starved for 4 h, and then fed with the above-mentioned diets. Samples were taken from each treatment group every 12 h; at each time point, 5 larvae were randomly dissected, and this was repeated three times in total. The larvae were rinsed with sterile water for 90 s, with 75 % ethanol for 90 s, and again with sterile water for 90 s. The guts were collected in 1.5-mL centrifuge tubes on a sterile dissection bench, and 1 mL of sterile water was added to each tube. After vortex mixing, the suspension was diluted 100-fold with sterile water, and 50 µL of the dilution was spread onto MYP agar plates [51]. The plates were incubated at 30 °C for 12 h, after which the growth of Bt was observed and photographed.

### In *Vitro* inhibitory effects of OV6 and Arg on Bt

In *vitro*: The Oxford cup agar diffusion assay was used to test the growth inhibitory effects of OV6 and Arg on Bt in *vitro* [51]. The OD_600_ of Bt and OV6 were 0.015 and 0.1 respectively, and the concentration of Arg was 5 mg/mL.

### OV6 genome sequencing

Activated OV6 bacterial cells were sent to Shanghai Majorbio Bio-pharm Technology Co., Ltd. for genome sequencing using the Illumina system. Sequence assembly software included SOAPdenovo2 v.2.04 and unicycler v.0.4.8, and the software for predicting coding sequences was Glimmer v.3.02 [52]. Finally, the KEGG database was used for functional annotation of the genome sequence [53], and Circos was used to create a genome overview to reveal the annotation [54]. The sequencing data was submitted to the NCBI: PRJNA1337595 (SRA).

### Construction of mutant strain OV6-Δ*argH*

Based on the OV6 genome sequence, its Arg synthesis pathway was analyzed (https://www.kegg.jp/pathway/map00220). The key gene *ArgH* in this synthesis pathway was cloned using primers ArgH-F/R and vector pESI-Blunt. The suicide plasmid pRE112 was used to construct the deletion mutant of the *ArgH* gene: 500 bp homologous arm sequences upstream and downstream of the *ArgH* gene were amplified with primers Up-F/R and Down-F/R, the KnaR resistance fragment was amplified from the pKD4 plasmid with primers KnaR-F/R, and they were fused together by overlap extension polymerase chain reaction (with the KnaR resistance fragment between the homologous arm sequences), and detected with primers Test1-F/R. The constructed recombinant plasmid pRE112-H digested with XbaI and XmaI was transformed into *E. coli* MC1061 and S17-1 λ pir competent cells successively and then transferred into the wild-type strain OV6 by electroporation to obtain OV6-pRE112-H, which was detected with primers Test2-F/R after transformation. Finally, the mutant strain with dual resistance to sucrose and kanamycin was screened on LB-sucrose solid medium (15 % sucrose), and the obtained mutant strain OV6-Δ*argH* was confirmed by detection with primers Test3-F/R and amplified sequence determination. Amplification and detection primers are shown in S2 Table.

### The ability of OV6 and OV6-Δ*argH* to synthesize Arg

Activated OV6 and OV6-Δ*argH* were cultured at 37 ℃ and 180 rpm for 48 h. 1.5 mL bacterial solution was centrifuged at 6000 rpm for 10 min at 4 ℃, 500 µL supernatant was taken into a 1.5 mL centrifuge tube, and the content of Arg synthesized by them was detected using the modified Sakaguchi reagent method [55].

### Detection of Arg content in *P. xylostella* gut

Each diet served as 1 treatment group, with 5 replicates per treatment group and 30 larvae per replicate. Late 3rd-instar *P. xylostella* larvae reared on the normal diet were randomly selected, starved for 4 h, fed specific diets for 48 h, and 10 larvae were randomly selected from each replicate for dissection to obtain gut tissue samples, where the gut tissue was fully ground. The Arg content in the obtained samples was then detected using the modified Sakaguchi reagent method [55].

### Reactive Oxygen Species Detection

Each diet served as 1 treatment group, with 3 replicates per treatment group and 70 larvae per replicate. Late 3rd-instar larvae fed normal diet were randomly selected, starved for 4 h, fed specific diets for 48 h, and 50 larvae randomly selected from each replicate of all treatments were dissected for gut tissue, followed by quantitative detection of reactive oxygen species using the Insect Reactive Oxygen Species (ROS) Detection Kit (Youxuan Biotechnology, Cat#YX-181519I). In addition, the production of ROS in gut epithelial cells was determined using the intracellular ROS-sensitive fluorescent dye dihydroethidium [56]: gut tissues were incubated in 5 μM dihydroethidium (Yeasen, Cas#38483-26-0) at room temperature in the dark for 30 min. Washed 3 times with 1×PBS (5 min each time), fixed with 4 % paraformaldehyde for 30 min, and washed again 3 times with 1×PBS. Scanned and photographed using a laser confocal microscope (TCS SP8, Leica, Germany).

### Sectioning and scanning electron microscopy of *P. xylostella* midgut tissue

Each diet served as 1 treatment group. Late 3rd-instar larvae (wild-type WT and mutant Δ*PxDES*) fed normal diet were randomly selected, starved for 4 h, fed specific diets for 48 h, and 10 larvae were randomly selected from all treatments for dissection of midgut tissue. Sectioning: the dissected gut was fixed in 4 % paraformaldehyde overnight, then paraffin-embedded, longitudinally sectioned, stained with hematoxylin-eosin, mounted, and observed for midgut cell and peritrophic membrane structure under a microscope.

Scanning electron microscopy: the midgut tissue was cut and attached to filter paper, rinsed 3 times with 1×PBS (8 min each time), then fixed with electron microscope fixative (2.5 % glutaraldehyde) at room temperature in the dark for 2 h, and stored at 4 ℃ overnight. The fixed samples were rinsed 3 times with 0.1M phosphate buffer PB (PH7.4) (15 min each time). Fixed with 1 % osmium tetroxide at room temperature in the dark for 1-2 h. Rinsed 3 times with 0.1M phosphate buffer PB (PH7.4) (15 min each time). The tissue was dehydrated with gradient ethanol of 30 %-50 %-70 %-80 %-90 %-95 %-100 %-100 % for 15 min each time, and isoamyl acetate for 15 min. The samples were dried in a critical point dryer (K850, Quorum), then attached to a conductive carbon film double-sided tape and placed on the sample stage of an ion sputter (MC1000, Hitachi) for gold sputtering for about 30 s. Finally, observed and photographed using a scanning electron microscope (SU8100, Hitachi).

### qRT-PCR detection

qRT-PCR of ceramide synthesis-related genes: each diet served as 1 treatment group. 3 replicates per treatment group, 30 larvae per replicate. Late 3rd-instar larvae (wild-type WT) fed normal diet were randomly selected, starved for 4 h, fed specific diets for 48 h, and 5 larvae were randomly selected from each replicate for total RNA extraction. The extraction of total RNA, reaction system and procedure of qRT-PCR, and calculation method of gene expression level all referred to the paper by Sun et al. [57]. The ribosomal protein RPL32 gene of *P. xylostella* was used as the internal reference gene for qRT-PCR. The primer sequences of the genes are shown in S3 Table; qRT-PCR of AMPs: this part refers to “qRT-PCR of ceramide synthesis-related genes”, with the difference that total RNA was extracted from the gut tissue of 20 larvae.

### Cloning of *PxSMPD1* and *PxDES* genes

This part refers to the paper by Sun et al. [57], with modifications to primers and Polymerase Chain Reaction (PCR) procedures. The amplification primers for the sequences of *PxSMPD1* (*Px003742*) and *PxDES* (*Px008081*) genes were Px003742-F/R and Px008081-F/R respectively. The PCR amplification procedure for *PxSMPD1* was: 95 ℃ for 3 min, 35 cycles at 95 ℃ for 15 s, 56 ℃ for 15 s, 72 ℃ for 90 s, and 72 ℃ for 5 min; the PCR amplification procedure for *PxDES* was: 95 ℃ for 3 min, 35 cycles at 95 ℃ for 15 s, 54 ℃ for 15 s, 72 ℃ for 45 s, and 72 ℃ for 5 min. Positive clones of *PxSMPD1* and *PxDES* genes were detected with universal primers M13F/R. The PCR amplification procedure for the detection of *PxSMPD1* positive clones was: 95 ℃ for 3 min, 35 cycles at 95 ℃ for 15 s, 55 ℃ for 15 s, 72 ℃ for 90 s, and 72 ℃ for 5 min; the PCR amplification procedure for the detection of *PxDES* positive clones was: 95 ℃ for 3 min, 35 cycles at 95 ℃ for 15 s, 55 ℃ for 15 s, 72 ℃ for 45 s, and 72 ℃ for 5 min. Primers are shown in S4 Table.

### Knockout of *PxSMPD1* and *PxDES* genes

This part refers to the paper by Sun et al. [57], with modifications to primers and PCR procedures for detecting mutations. The upstream primer for the *PxSMPD1* knockout target was Px003742-gRNA-F, the upstream primer for the *PxDES* knockout target was Px008081-gRNA-F, and the universal downstream primer for the two gene knockout targets was gRNA-comR. The detection primers for *PxSMPD1* mutations were Px003742-T-F/R, and the detection primers for *PxDES* mutations were Px008081-T-F/R. The PCR procedures for mutation detection of both genes were: 95 ℃ for 3 min, 35 cycles at 95 ℃ for 15 s, 54 ℃ for 15 s, 72 ℃ for 1 min, and 72 ℃ for 5 min. Primers are shown in S4 Table.

### Retrieval of key enzymes in the ceramide synthesis pathway across eukaryotes

The key enzymes involved in ceramide synthesis within the sphingolipid metabolic pathway—namely, serine palmitoyltransferase (SPT), 3-ketodihydrosphingosine reductase (KDSR), ceramide synthase (CERS), dihydroceramide desaturase (DES), sphingomyelin phosphodiesterase (SMPD), and glucocerebrosidase (GBA)—were searched and compared against the genomes of six eukaryotic species, including *D. melanogaster*, *Mus musculus*, *Danio rerio*, *Homo sapiens*, *Sus scrofa*, and *Gallus gallus*, using the NCBI (National Center for Biotechnology Information) database. The presence of the corresponding key enzyme genes in a given host was taken as evidence that the host possesses these enzymes.

### Omics analysis of *P. xylostella* under Arg treatment

Each diet served as 1 treatment, with 3 replicates for transcriptome and 6 replicates for metabolome, 10 larvae per replicate. Late 3rd-instar larvae fed the normal diet were randomly selected, starved for 4 h, fed specific diets for 48 h, rinsed on the body surface with sterile 1×PBS in a sterile dissecting table, quick-frozen in liquid nitrogen for 0.5 hours, stored in a –80 ℃ refrigerator, and sent to Shanghai Majorbio Bio-pharm Technology Co., Ltd. For transcriptome and metabolome sequencing after collection. The sequencing data was submitted to the China National Center for Bioinformation (CNCB): PRJCA049482 and PRJCA049624.

#### Transcriptome analysis

The total RNA as a template, the mRNA library was constructed with the TruSeq™ mRNA Kit (Illumina, San Diego, CA) and then sequenced on the DNBSEQ-T7 platform. The fastp software [58] was used to filter and quality control the raw data to obtain clean data. The Hisat2 software [59] was used to align clean data with the *P. xylostella* reference genome [25] to obtain mapped reads. The StringTie software [60] was used to assemble the mapped reads and functionally annotate the transcripts. The RSEM software [61] was used for quantitative analysis of gene expression levels. The DESeq2 software [62] was used for differential gene expression analysis between groups, with FC (Fold Change) ≥ 2 and P adjust < 0.05 as screening criteria. The KEGG database was used for enrichment analysis of differential genes.

#### Metabolome analysis

The UHPLC-Triple TOF system (AB SCIEX) was used for metabolic profile detection. The chromatographic column was ACQUITY UPLC HSS T3 (100 mm × 2.1 mm, 1.8 µm; Waters, USA); 95 % water + 5 % acetonitrile (containing 0.1 % formic acid) was used as mobile phase A, and 47.5% acetonitrile + 47.5 % isopropanol + 5 % water (containing 0.1 % formic acid) as mobile phase B; the flow rate was 0.4 mL/min, and the column temperature was 40 ℃. The sample volume was 10 μL. Elution was performed according to the following gradient: 0-0.5 min, 100 %A; 0.5-2.5min, 100-75 %A; 2.5-9min, 75-0 %A; 9-13min, 0 %A; 13-13.1min, 0-100 %A; 13.1-16min, 100 %A. The samples were collected in positive and negative ion scanning modes of the Electrospray Ionization (ESI) source: ion source temperature: 550 ℃, ion source voltage: 5000 V in positive ion mode, –4000 V in negative ion mode. Mass spectrometry scanning range m/z: 50-1000.

The ropls (R packages) (Version1.6.2) software was used for Principal Component Analysis (PCA) and Partial Least Squares Discriminant Analysis (PLS-DA) of the obtained data. Then, with VIP value >1 of the first principal component of the PLS-DA model, P value <0.05, and FC (Fold Change) >2 or <0.83 as screening criteria, differential metabolites between groups were screened out. The KEGG database was used for enrichment analysis of differential metabolites.

#### Integrated transcriptome and metabolome analysis

The differential genes obtained from transcriptome analysis and differential metabolites obtained from metabolome analysis were subjected to KEGG pathway association analysis to obtain the common pathway information of the two and clarify the main biochemical pathways and signal transduction pathways jointly involved by differentially expressed genes and differential metabolites.

### Transcriptome analysis of *P. xylostella* under ceramide treatment

Each diet served as one treatment group, with 3 replicates per treatment group and 30 larvae per replicate. Late 3rd-instar larvae (wild-type WT and mutant Δ*PxDES*) fed normal diet were randomly selected, starved for 4 h, fed specific diets for 48 h, dissected for larval guts in a sterile dissecting table, quick-frozen in liquid nitrogen for 0.5 h, stored in a –80 ℃ refrigerator, and sent to Novogene Bioinformatics Technology Co., Ltd. for transcriptome sequencing after collection. The sequencing and analysis process refers to “Omics analysis of *P. xylostella* under Arg treatment”. The sequencing data was submitted to the China National Center for Bioinformation (CNCB): PRJCA049624.

### 16S rRNA sequencing analysis of *P. xylostella* under ceramide treatment

Each diet was designated as 1 treatment group, with 3 biological replicates per group and 50 larvae per replicate. Late 3rd-instar larvae (wild-type WT and mutant Δ*PxDES*) reared on the normal diet were randomly selected, starved for 4 h and then fed the specific diets for 48 h. The larval midguts were dissected on a sterile dissecting bench, snap-frozen in liquid nitrogen for 0.5 h and subsequently stored at –80 °C. After sample collection, all specimens were sent to Novogene Bioinformatics Technology Co., Ltd. for 16S rRNA sequencing. The sequencing data was submitted to the China National Center for Bioinformation (CNCB): PRJCA049482.

PCR amplification was performed with bacterial genomic DNA as the template, using primers 338F (5′-ACTCCTACGGGAGGCAGCAG-3′) and 806R (5′-GGACTACHVGGGTWTCTAAT-3′) that target the V3-V4 hypervariable regions of the bacterial 16S rRNA gene (where H, V and W represent different degenerate bases) [63]. The 25 μL PCR reaction system consisted of 5 μL of 5× reaction buffer, 5 μL of 5× GC buffer, 2 μL of 2.5 mM dNTPs, 1 μL of 10 μM 338F, 1 μL of 10 μM 806R, 2 μL of DNA template, 8.75 μL of ddH₂O and 0.25 μL of Q5 DNA Polymerase. The PCR program was as follows: initial denaturation at 98 °C for 1 min, followed by 30 cycles of denaturation at 98 °C for 10 s, annealing at 50 °C for 30 s and extension at 72 °C for 30 s, with a final extension at 72 °C for 5 min. The PCR products were purified with the Vazyme VAHTS™ DNA Clean Beads (N411-01, Vazyme Biotech Co., Ltd), and fluorescent quantification of the purified products was conducted with the Quant-iT PicoGreen dsDNA Assay Kit (P11496, Thermo Fisher Scientific (China) Co., Ltd) with a Microplate Reader (FLx800, BioTek).

Sequencing libraries were constructed from the purified products with the TruSeq Nano DNA LT Library Prep Kit, and paired-end sequencing was performed on the Illumina NovaSeq 6000 platform. Sample-specific data were demultiplexed from the raw sequencing data based on barcode and PCR primer sequences. After trimming the barcode and primer sequences, the reads of each sample were assembled using FLASH (Version 1.2.11) [64] to generate raw tag data (Raw Tags). The Cutadapt software was used to match reverse primer sequences and trim the remaining fragments. High-quality tag data (Clean Tags) were obtained by filtering the assembled Raw Tags with fastp (Version 0.23.1) [65]. Chimeric sequences in the tag sequences were detected by alignment against taxonomic annotation databases—the SILVA database (https://www.arb-silva.de/) for 16S/18S rRNA and the UNITE database (https://unite.ut.ee/) for ITS—and finally removed to yield the final effective data (Effective Tags) [66]. Effective Tags were denoised using QIIME2 (Version QIIME2-202202) to obtain the final Amplicon Sequence Variants (ASVs) and feature tables [67], and taxonomic annotation was performed with QIIME2. The Shannon index (http://scikitbio.org/docs/latest/generated/skbio.diversity.alpha.shannon.html#skbio.diversity.alpha.shannon) was subsequently used to calculate microbial community diversity.

### Ceramide-mediated in *vitro* inhibitory effects on bacteria

The Oxford cup agar diffusion method [51] was used to determine the in *vitro* growth inhibitory effects of ceramide on Bt, Sm, Sa, OV6, *Enterobacter* sp., *Pseudomonas parafulva*, *Kluyvera intermedia*, *Exiguobacterium acetylicum*, *Microbacterium arborescens*, *Carnobacterium gallinarum* and *Bacillus cereus*, with all bacterial strains adjusted to an OD₆₀₀ of 0.02 and the ceramide concentration set at 5 mg/mL.

The minimum inhibitory concentration (MIC) of ceramide against Bt, Sm and Sa were determined via the broth microdilution method [68].

The bacteriostatic curve assay: Bt, Sm and Sa were activated and inoculated into liquid LB medium to an OD₆₀₀ of 0.5-0.6, then diluted with 1× PBS to a concentration of 1×10⁵ CFU/mL for subsequent use. Ceramide at different concentrations was co-cultured with the bacteria in 96-well plates (100 μL each, with a total volume of 200 μL), resulting in final ceramide concentrations of 1/2MIC, MIC and 2MIC, and an initial bacterial concentration of 1×10⁵ CFU/mL. Kanamycin (final concentration: 50 μg/mL) was used as the positive control, and 1× PBS as the negative control. After co-cultivation, 100 μL samples were collected at 0 h, 1 h, 2 h, 4 h, 6 h, 8 h, 12 h and 24 h, spread on solid LB medium, and colony-forming units (CFUs) were counted after 12 h of incubation. Three biological replicates were set for each treatment, with three technical replicates (plates) per biological replicate.

Scanning Electron Microscopy (SEM) of Bacteria: Bt, Sm and Sa were activated and inoculated into liquid LB medium to an OD₆₀₀ of 0.5–0.6, then treated with ceramide at final concentrations of 1/2MIC, MIC and 2MIC and further cultured for 30 h. Bacterial cells were harvested by centrifugation at 6000 rpm for 10 min at 4 °C, gently washed with 100 μL of 1× PBS and re-centrifuged at 6000 rpm for 10 min at 4 °C. The bacterial pellet was resuspended in 150 μL of electron microscopy fixative (2.5 % glutaraldehyde) by gentle pipetting, fixed at room temperature in the dark for 30 min, and then stored at 4 °C for subsequent use. The post-fixation processing and imaging of the samples followed the same protocol as described in the section of “Scanning electron microscopy of *P. xylostella* midgut tissue”.

Transmission electron microscopy (TEM) of Bt: Bt cells were fixed according to the SEM protocol, then centrifuged and the supernatant discarded. The pellet was washed 3 times (3 min each) with 0.1 M phosphate buffer (PB, pH 7.4), then picked up with forceps and suspended in 1% agarose before solidification for embedding. The samples were fixed with 1% osmium tetroxide at room temperature in the dark for 1–2 h, and washed 3 times (15 min each) with 0.1 M PB (pH 7.4). Gradient dehydration was performed with 30%, 50%, 70%, 80%, 95% and 100% ethanol (twice for 100% ethanol) for 20 min per gradient, followed by two rinses with 100% acetone for 15 min each. Pour the pure EMBed 812 into the embedding models and insert the tissues into the pure EMBed 812, and then keep in 37 ℃ oven overnight. The embedding models with resin and samples were moved into 60℃ oven to polymerize for 48 h. And then the resin blocks were taken out from the embedding models for standby application at room temperature. The resin blocks were cut to 60-80 nm thin on the ultra-microtome, and the tissues were fished out onto the 150 meshes cuprum grids with formvar film. 2% uranium acetate saturated alcohol solution avoid light staining for 8 min, rinsed in 70 % ethanol for 3 times and then rinsed in ultra-pure water for 3 times. 2.6 % Lead citrate avoid CO_2_ staining for 8 min, and then rinsed with ultra-pure water for 3 times. After dried by the filer paper, the cuprum grids were put into the grids board and dried overnight at room temperature. Imaging was finally performed using a transmission electron microscope (HT7800, Hitachi).

### Quantification and statistical analysis

All statistical analyses were performed with SPSS Statistics (Version 26.0). The Shapiro-Wilk test was used to assess the normality of data distribution. For statistical comparisons, independent-samples t-tests (for two groups) or one-way analysis of variance (ANOVA, for multiple groups) were applied to normally distributed data, while the Mann-Whitney U test (for two groups) or Kruskal-Wallis test (for multiple groups) was used for non-normally distributed data. Survival curves were analyzed using the Mantel-Cox test in GraphPad Prism (Version 9.0). Statistical significance was defined as follows: *p* < 0.05(*), *p <* 0.01(**), *p <* 0.001(***), *p* < 0.0001 (****); “ns” indicated no significant difference. The p-value < 0.05 was considered statistically significant.

## Acknowledgements

We thank our colleagues at the College of Plant Protection, Fujian Agriculture and Forestry University, the State Key Laboratory of Agricultural and Forestry Biosecurity, and the Institute of Applied Ecology for their support and discussions. We acknowledge the technical assistance provided by Shanghai Majorbio Bio-pharm Technology Co., Ltd. and Novogene Bioinformatics Technology Co., Ltd. in sequencing services. This research was funded by the National Natural Science Foundation of China (No. 31871968), the Natural Science Foundation of Fujian Province (2022J01126), and the Science and Technology Innovation Fund of Fujian Agriculture and Forestry University (KFB23024A).

## Author contributions

S.H. and M.L. designed the experiments and wrote the initial draft of the manuscript. G.M.G., L.V., M.Y., and X.X. conceptualized the project and supervised the research. All authors participated in experimental design, data analysis, and manuscript revision.

## Data availability

The transcriptomic, metabolomic, and 16S rRNA sequencing data generated in this study have been deposited in the China National Center for Bioinformation (CNCB) under accession numbers PRJCA049482 (transcriptome and 16S rRNA) and PRJCA049624 (metabolome). The genome sequence of *Raoultella ornithinolytica* (OV6) has been deposited in the National Center for Biotechnology Information (NCBI) Sequence Read Archive under accession number PRJNA1337595. All other data needed to evaluate the conclusions of this study are present in the paper or the extended data.

## Disclaimer

During manuscript preparation, the authors used Microsoft Copilot and Deepseek to assist with language editing, text refinement, and stylistic improvements. All scientific content, interpretations, analyses, and conclusions were developed, verified, and approved by the authors, who take full responsibility for the manuscript.

## References

1. Chen RC, Zou J, Chen JW, Zhong X, Kang R, Tang DL. Pattern recognition receptors: function, regulation and therapeutic potential. Signal Transduct. Target. Ther. 2025; 10(1): 216. 10.1038/s41392-025-02264-1 PMID: 40640149

2. Medzhitov R, Janeway CA Jr. Decoding the patterns of self and nonself by the innate immune system. Science. 2002; 296(5566): 298–300. 10.1126/science.1068883 PMID: 11951031

3. Cavaillon JM. The historical milestones in the understanding of leukocyte biology initiated by Elie Metchnikoff. J. Leukoc. Biol. 2011; 90(3): 413–424. 10.1189/jlb.0211094 PMID: 21628329

4. Ochoa MC, Minute L, Rodriguez I, Garasa S, Perez-Ruiz E, Inogés S, et al. Antibody-dependent cell cytotoxicity: immunotherapy strategies enhancing effector NK cells. Immunol. Cell Biol. 2017; 95(4): 347–355. 10.1038/icb.2017.6. PMID: 28138156

5. Mathis D, Shoelson SE. Immunometabolism: an emerging frontier. Nat. Rev. Immunol. 2011; 11(2): 81. 10.1038/nri2922 PMID: 21469396

6. Malla S, Shahreen N, Saha R. Immunometabolism at the crossroads of infecti on: mechanistic and systems-level perspectives from host and pathogen. Immu nometabolism (Cobham).2025; 7(4): e00069. 10.1097/IN9.0000000000000069 PMID: 41159060

7. Lv YY, Li ZS, Liu ST, Zhou ZK, Song JL, Ba YH, et al. Metabolic checkpoints in immune cell reprogramming: rewiring immunometabolism for cancer therapy. Mol. Cancer. 2025; 24(1): 210. 10.1186/s12943-025-02407-6 PMID: 40753443

8. Sun LZ, Yang XF, Yuan ZY, Wang H. Metabolic reprogramming in immune response and tissue inflammation. Arterioscler. Thromb. Vasc. Biol. 2020; 40(9): 1990–2001. 10.1161/ATVBAHA.120.314037 PMID: 32698683

9. Michelucci A, Cordes T, Ghelfi J, Pailot A, Reiling N, Goldmann O, et al. Immune-responsive gene 1 protein links metabolism to immunity by catalyzing itaconic acid production. Proc. Natl. Acad. Sci. 2013; 110(19): 7820–7825. 10.1073/pnas.1218599110 PMID: 23610393

10. Chen LL, Morcelle C, Cheng ZL, Chen XF, Xu YP, Gao YJ, et al. Itaconate inhibits TET DNA dioxygenases to dampen inflammatory responses. Nat. Cell Biol. 2022; 24(3): 353–363. 10.1038/s41556-022-00853-8 PMID: 35256775

11. Chen C, Li XJ. The cell autonomous and non-autonomous roles of itaconate i n immune response. Cell Insight. 2024; 4(1): 100224. 10.1016/j.cellin.2024.100224 PMID: 39877254

12. Jiang XC, Li ZQ. Sphingolipids and cholesterol. Adv. Exp. Med. Biol. 2022; 1372: 1–14. 10.1007/978-981-19-0394-6_1 PMID: 35503170

13. Simons K, Sampaio JL. Membrane organization and lipid rafts. Cold Spring Harb. Perspect. Biol. 2011; 3(10): a004697. 10.1101/cshperspect.a004697 PMID: 21628426

14. Schmieder SS, Tatituri R, Anderson M, Kelly K, Lencer WI. Structural basis for acyl chain control over glycosphingolipid sorting and vesicular trafficking. Cell Rep. 2022; 40(2): 111063. 10.1016/j.celrep.2022.111063 PMID: 35830800

15. Obeid LM, Linardic CM, Karolak LA, Hannun YA. Programmed cell death in duced by ceramide. Science. 1993; 259(5102): 1769–1771. 10.1126/science.8456305 PMID: 8456305

16. Jiang W, Ogretmen B. Autophagy paradox and ceramide. Biochim. Biophys. Acta. 2014; 1841(5): 783–792. 10.1016/j.bbalip.2013.09.005 PMID: 24055889

17. Liu FY, Li X, Lu CW, Bai AP, Bielawski J, Bielawska A, et al. Ceramide activates lysosomal cathepsin B and cathepsin D to attenuate autophagy and induces ER stress to suppress myeloid-derived suppressor cells. Oncotarget 2016; 7(51): 83907–83925. 10.18632/oncotarget.13438 PMID: 27880732

18. Summers SA, Chaurasia B, Holland WL. Metabolic messengers: ceramides. Nat. Metab. 2019; 1(11): 1051–1058. 10.1038/s42255-019-0134-8 PMID: 32694860

19. Becam J, Walter T, Burgert A, Schlegel J, Sauer M, Seibel J, et al. Antibacterial activity of ceramide and ceramide analogs against pathogenic *Neisseria*. Sci. Rep. 2017; 7(1): 17627. 10.1038/s41598-017-18071-w PMID: 29247204

20. Nocedo-Mena D, Arrasate S, Garza-González E, Rivas-Galindo VM, Romo-Mancillas A, Munteanu CR, et al. Molecular docking, SAR analysis and biophysical approaches in the study of the antibacterial activity of ceramides isolated from Cissus incisa. Bioorg. Chem. 2021; 109: 104745. 10.1016/j.bioorg.2021.104745 PMID: 33640629

21. Paudel S, Wu G, Wang X. Amino acids in cell signaling: regulation and function. Adv. Exp. Med. Biol. 2021; 1332: 17–33. 10.1007/978-3-030-74180-8_2 PMID: 34251636

22. Li W, Jin M, Li HY, Zhong RY, Lin ZH, Shen M, et al. Amino acid and cholesterol metabolism in innate immunity. Mol. Cell. 2025; 85(20): 3760–3778. 10.1016/j.molcel.2025.09.019 PMID: 41106367.

23. Kelly B, Pearce EL. Amino assets: how amino acids support immunity. Cell Metab. 2020; 32(2): 154–175. 10.1016/j.cmet.2020.06.010 PMID: 32649859

24. Pan J, Lin Y, Liu X, Zhang X, Liang T, Bai X. Harnessing amino acid pathways to influence myeloid cell function in tumor immunity. Mol. Med. 2025; 31(1): 44. 10.1186/s10020-025-01099-4 PMID: 39905317

25. You MS, Yue Z, He WY, Yang XH, Yang G, Xie M, et al. A heterozygous moth genome provides insights into herbivory and detoxification. Nat. Genet. 2013; 45(2): 220–225. 10.1038/ng.2524 PMID: 23313953

26. Furlong MJ, Wright DJ, Dosdall LM. Diamondback moth ecology and management: problems, progress and prospects. Annu. Rev. Entomol. 2013; 58: 517–541. 10.1146/annurev-ento-120811-153605 PMID: 23020617

27. Chen YT, Liu ZX, Régnière J, Vasseur L, Lin J, Huang SG, et al. Large-scale genome-wide study reveals climate adaptive variability in a cosmopolitan pest. Nat. Commun. 2021; 12(1): 7206. 10.1038/s41467-021-27510-2 PMID: 34893609

28. Hannun Y.A, Obeid LM. Principles of bioactive lipid signalling: lessons from sphingolipids. Nat. Rev. Mol. Cell Biol. 2008; 9(2): 139–150. 10.1038/nrm2329 PMID: 18216770

29. Velazquez FN, Hernandez-Corbacho M, Trayssac M, Stith JL, Bonica J, Jean B, et al. Bioactive sphingolipids: Advancements and contributions from the la boratory of Dr. Lina M. Obeid. Cell. Signal. 2021; 79: 109875. 10.1016/j.cellsig.2020.109875 PMID: 33290840

30. Ryu JH, Ha EM, Lee WJ. Innate immunity and gut-microbe mutualism in *Dr osophila*. Dev. Comp. Immunol. 2010; 34(4): 369–376. 10.1016/j.dci.2009.11.010 PMID: 19958789

31. Hanson MA, Grollmus L, Lemaitre B. Ecology-relevant bacteria drive the evolution of host antimicrobial peptides in *Drosophila*. Science. 2023; 381(6655), eadg5725. 10.1126/science.adg5725 PMID: 37471548

32. Guo LZ, Tang JB, Tang M, Luo SQ, Zhou X. Reactive oxygen species are regulated by immune deficiency and toll pathways in determining the host specificity of honeybee gut bacteria. Proc. Natl. Acad. Sci. 2023; 120(33): e2219634120. 10.1073/pnas.2219634120 PMID: 37556501

33. Yang YJ, Liu XG, Xu HX, Liu YH, Lu ZX. Effects of host plant and insect generation on shaping of the gut microbiota in the Rice Leaffolder, *Cnaphaloc rocis medinalis*. Front. Microbiol. 2022; 13: 824224. 10.3389/fmicb.2022.824224 PMID: 35479615

34. Wang AL, Yao ZC, Zheng WW, Zhang HY. Bacterial communities in the gut and reproductive organs of *Bactrocera minax* (Diptera: Tephritidae) based on 4 54 pyrosequencing. PLoS One. 2014; 9(9): e106988. 10.1371/journal.pone.0106988 PMID: 25215866

35. Han SC, Ai QQ, Xia XF. Potential source and transmission pathway of gut bacteria in the Diamondback Moth, *Plutella xylostella*. Insects. 2023; 14(6): 504. 10.3390/insects14060504 PMID: 37367320

36. Chen JL, Duan YW, Zhou YY, Yang Q. Squeeze pumping of lipids and insec ticides by ABCH transporter. Cell. 2025; 188(4): 944–957. 10.1016/j.cell.2024.11.033 PMID: 39721587

37. Silhavy TJ, Kahne D, Walker S. The bacterial cell envelope. Cold Spring Harb Perspect Biol. 2010; 2(5): a000414. 10.1101/cshperspect.a000414 PMID: 20452953

38. Thia JA, Dorai APS, Hoffmann AA. Symbiotic bacteria and pest control: plant toxins, chemical pesticides, and fungal entomopathogens. Trends Microbiol. 2025; 33(9): 976–988. 10.1016/j.tim.2025.04.007 PMID: 40320312

39. Bai YR, Liu WX, Liu ZY, Ding DD, Jin HY, Xiao SY, et al. Multi-omics analysis of glycolytic reprogramming and ROS dynamics in host-specific responses to *Salmonella Typhi* infection in mice. J. Adv. Res. 2026; 80: 435–450. 10.1016/j.jare.2025.05.027 PMID: 40393644

40. Hajjar R, Ambaraghassi G, Sebajang H, Schwenter F, Su SH. Raoultella ornithinolytica: Emergence and resistance. Infect. Drug Resist. 2020; 13: 1091–1104. 10.2147/IDR.S191387 PMID: 32346300

41. Indiragandhi P, Anandham R, Madhaiyan M, Poonguzhali S, Kim GH, Saravanan VS, et al. Cultivable bacteria associated with larval gut of prothiofos-resistant, prothiofos-susceptible and field-caught populations of diamondback moth, *Plutella xylostella* and their potential for antagonism towards entomopathogenic fungi and host insect nutrition. J. Appl. Microbiol. 2007; 103(6): 2664–2675. 10.1111/j.1365-2672.2007.03506.x PMID: 17973916

42. Liao WY, Lv ZH, Mo YF, Yu SW, Zhao YF, Zhang SS, et al. Battle between gut bacteria, immune system, and Cry1Ac toxin in *Plutella xylostella*. J. Agric. Food Chem. 2025; 73(18): 10896–10906. 10.1021/acs.jafc.5c01838 PMID: 40271618

43. Sun XM, Wang YH, Yuan F, Zhang YN, Kang X, Sun J, et al. Gut symbiont-derived sphingosine modulates vector competence in *Aedes mosquitoes*. Nat. Commun. 2024; 15(1): 8221. 10.1038/s41467-024-52566-1 PMID: 39300135

44. Hammermann R, Brunn G, Racké K. Analysis of the genomic organization of the human cationic amino acid transporters CAT-1, CAT-2 and CAT-4. Amino Acids. 2001; 21(2): 211–219. 10.1007/s007260170029 PMID: 11665818

45. Verrey F, Closs EI, Wagner CA, Palacin M, Endou H, Kanai Y. CATs and HATs: the SLC7 family of amino acid transporters. Pflugers Arch. 2004; 447(5): 532–542. 10.1007/s00424-003-1086-z PMID: 14770310

46. Ge YS, Li F, He Y, Cao Y, Guo WJ, Hu GQ, et al. L-arginine stimulates the proliferation of mouse mammary epithelial cells and the development of mammary gland in pubertal mice by activating the GPRC6A/PI3K/AKT/mTOR signalling pathway. J. Anim. Physiol. Anim. Nutr. (Berl). 2022; 106(6): 1383–1395. 10.1111/jpn.13730 PMID: 35616019

47. Wang SY, Tsun ZY, Wolfson RL, Shen K, Wyant GA, Plovanich ME, et al. Lysosomal amino acid transporter SLC38A9 signals arginine sufficiency to mT ORC1. Science. 2015; 347(6218): 188–194. 10.1126/science.1257132 PMID: 25567906

48. Lee G, Zheng YX, Cho S, Jang C, England C, Dempsey JM, et al. Post-tran scriptional regulation of *de novo* lipogenesis by mTORC1-S6K1-SRPK2 signali ng. Cell. 2017; 171(7): 1545–1558. 10.1016/j.cell.2017.10.037 PM ID: 29153836

49. Lin JH, Yu XQ, Wang Q, Tao XP, Li JY, Zhang SS, et al. Immune response s to Bacillus thuringiensis in the midgut of the diamondback moth, *Plutella x ylostella*. Dev. Comp. Immunol. 2020; 107: 103661. 10.1016/j.dci.2020.103661 PMID: 32097696

50. Xu XF, Yu JP. Determination of 17 free amino acids in human serum by AQC pre-column derivatization-UPLC. Int. J. Lab. Med. 2014; 35(5): 596–597. 10.3969/j.issn.1673-4130.2014.05.038 CNKI: 1673-4130 (2014) 05-0596-02

51. Tao XP, Jia YH, Sun Y, Han SC, Xia XF. Gut bacteria reduce the Bt susceptibility in the diamondback moth, *Plutella xylostella* (Lepidoptera: Plutellidae), by competing for niche and protecting the inner wall of gut. Acta. Entomol. Sin. 2022; 65(12): 1645–1657. 10.16380/j.kcxb.2022.12.010 CNKI: 0454-6296 (2022) 12-1645-13

52. Delcher, AL, Bratke KA, Powers EC, Salzberg SL. Identifying bacterial genes and endosymbiont DNA with Glimmer. Bioinformatics. 2007; 23(6): 673–679. 10.1093/bioinformatics/btm009 PMID: 17237039

53. Kanehisa M, Furumichi M, Tanabe M, Sato Y, Morishima K. KEGG: new perspectives on genomes, pathways, diseases and drugs. Nucleic Acids Res. 2017; 45(D1): D353–D361. 10.1093/nar/gkw1092 PMID: 27899662

54. Krzywinski M, Schein J, Birol I, Connors J, Gascoyne R, Horsman D, et al. Circos: an information aesthetic for comparative genomics. Genome Res. 2009; 19(9): 1639–1645. 10.1101/gr.092759.109 PMID: 19541911

55. Hao G, Qian H. Study on quantitative detection methods of L-Arginine. Sci. Technol. Food Ind. 2005; 26(2): 184–186. 10.13386/j.issn1002-0306.2005.02.058 CNKI: 1002-0306 (2005)02-0184-03

56. Wei G, Lai YL, Wang GD, Chen H, Li F, Wang SB. Insect pathogenic fungus interacts with the gut microbiota to accelerate mosquito mortality. Proc. Natl. Acad. Sci. U.S.A. 2017; 114(23): 5994–5999. 10.1073/pnas.1703546114 PMID: 28533370

57. Sun Y, Yuan J, Han SC, Wang QP, Akhtar MR, Xia XF. PxSpätzle3 regulates the Toll pathway to affect *Bacillus thuringiensis* susceptibility of *Plutella xyl ostella*. J. Agric. Food Chem. 2025; 73(9): 5129–5139. 10.1021/acs.jafc.4c11380 PMID: 39965920

58. Chen SF, Zhou YQ, Chen YR, Gu J. Fastp: an ultra-fast all-in-one FASTQ pr eprocessor. Bioinformatics. 2018; 34(17): i884–i890. 10.1093/bioinformatics/bty560 PMID: 30423086

59. Kim D, Langmead B, Salzberg SL. HISAT: a fast spliced aligner with low m emory requirements. Nat. Methods. 2015; 12(4): 357–360. 10.1038/nmeth.3317 PMID: 25751142

60. Pertea M, Pertea GM, Antonescu CM, Chang T, Mendell JT, Salzberg SL. StringTie enables improved reconstruction of a transcriptome from RNA-seq reads. Nat. Biotechnol. 2015; 33(3): 290–295. 10.1038/nbt.3122 PMID: 25690850

61. Li B, Dewey CN. RSEM: accurate transcript quantification from RNA-Seq data with or without a reference genome. BMC Bioinformatics. 2011; 12: 323. 10.1186/1471-2105-12-323 PMID: 21816040

62. Love MI, Huber W, Anders S. Moderated estimation of fold change and dispersion for RNA-seq data with DESeq2. Genome Biol. 2014; 15(12): 550. 10.1186/s13059-014-0550-8 PMID: 25516281

63. Dennis KL, Wang YW, Blatner NR, Wang SY, Saadalla A, Trudeau E, et al. Adenomatous polyps are driven by microbe-instigated focal inflammation and are controlled by IL-10-producing T cells. Cancer Res. 2013; 73(19): 5905–5913. 10.1158/0008-5472.CAN-13-1511 PMID: 23955389

64. Magoč T, Salzberg SL. FLASH: fast length adjustment of short reads to impr ove genome assemblies. Bioinformatics. 2011; 27(21): 2957–2963. 10.1093/bioinformatics/btr507 PMID: 21903629

65. Bokulich NA, Subramanian S, Faith J, Gevers D, Gordon JI, Knight R, et al. Quality-filtering vastly improves diversity estimates from Illumina amplicon se quencing. Nat. Methods. 2013; 10(1): 57–59. 10.1038/nmeth.2276 PMID: 23202435

66. Edgar RC, Haas BJ, Clemente JC, Quince C, Knight R. UCHIME improves sensitivity and speed of chimera detection. Bioinformatics. 2011; 27(16): 2194–2200. 10.1093/bioinformatics/btr381 PMID: 21700674

67. Wang YY, Guo H, Gao XG, Wang JH. The intratumor microbiota signatures associate with subtype, tumor stage, and survival status of esophageal carcinoma. Front. Oncol. 2021; 11: 754788. 10.3389/fonc.2021.754788 PMID: 34778069

68. Arab Y, Şahin B, Ceylan O, Zellagui A, Olmez OT, Kucukaydin S, et al. As sessment of in vitro activities and chemical profiling of Senecio hoggariensis growing in Algerian Sahara. Biodiversitas. 2022; 23(7): 3498–3506. 10.13057/biodiv/d230724 E-ISSN: 2085-4722

