## Supplementary material for "Arginine activates a conserved sphingolipid defense axis against infection": Supporting information.pdf

### This file includes:

Supporting Figures 1-7
Supporting Tables 1-4

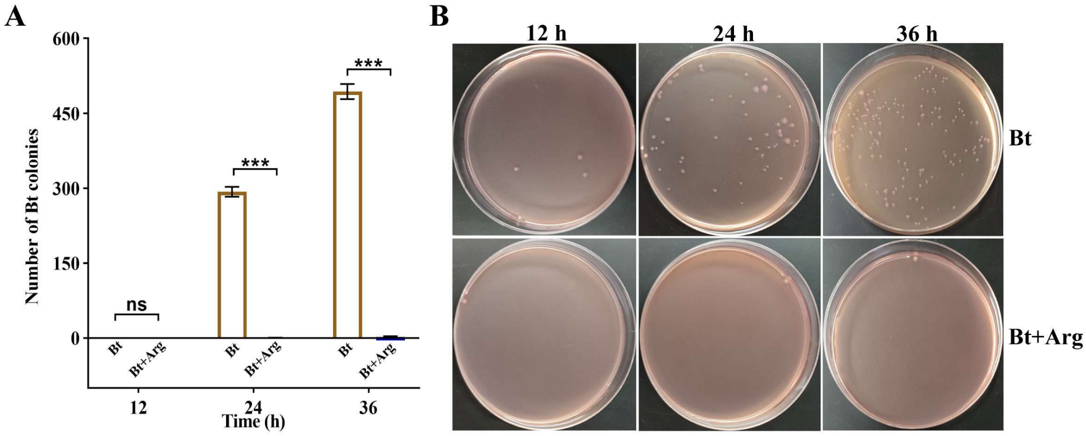


**S1 Fig. Arginine inhibits Bt growth in the gut of *P. xylostella.*** (**A**) Arginine suppresses Bt growth in the gut as determined by bacterial colony counting; n = 3 biological replicates. (**B**) Arginine suppresses Bt growth in the gut (representative images of Bt growth). Abbreviations of the full names are used in the x-axis, with the details provided in the Materials and Methods section. For (**A**), the independent samples t test was used; asterisks indicate $P < 0.001$ (***), "ns" denotes no significant difference. $P < 0.05$ is considered statistically significant.

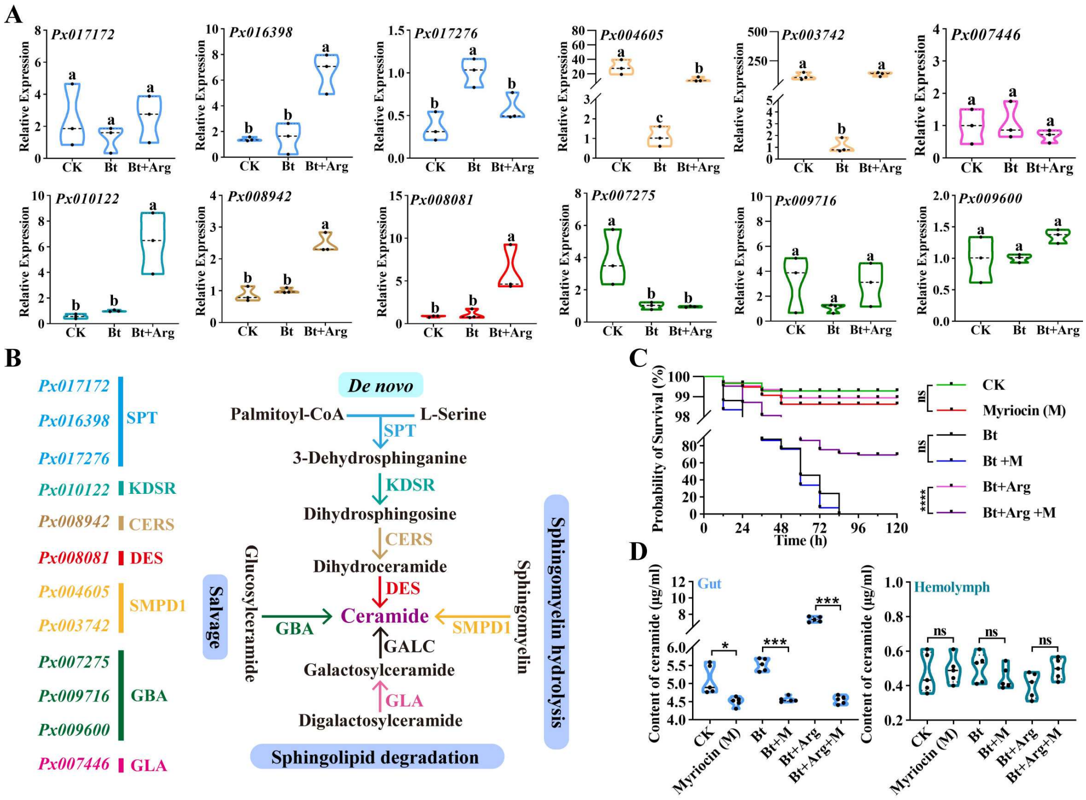


**S2 Fig. Arginine activates the ceramide synthesis pathway within the sphingolipid metabolic pathway.** (**A**) Arginine upregulates the expression of ceramide synthesis related genes in the sphingolipid metabolic pathway of *P. xylostella*; n = 3 biological replicates. (**B**) Schematic of the ceramide synthesis pathway in the sphingolipid metabolic pathway of *P. xylostella*. (**C**) The SPT inhibitor Myriocin attenuates the ability of arginine to mediate host defense against Bt in *P. xylostella*; n = 3 biological replicates. (**D**) The SPT inhibitor Myriocin suppresses the arginine induced increase in gut ceramide levels in *P. xylostella*; n = 5 biological replicates. The genes encoding serine palmitoyltransferase (SPT) are *Px017172*, *Px016398*, and *Px017276*, 3 ketodihydrosphingosine reductase (KDSR) is *Px010122*, ceramide synthase (CERS) is *Px008942*, dihydroceramide desaturase (DES) is *Px008081*, sphingomyelin phosphodiesterase 1 (SMPD1) are *Px004605* and *Px003742*, glucosylceramidase (GBA) are *Px007275*, *Px009716*, and *Px009600*, and alpha galactosidase A (GLA) is *Px007446*. Abbreviations of the full names are used in the x-axis, with the details provided in the Materials and Methods section. For (**A**), one-way ANOVA was performed, and different lowercase letters indicate significant differences ($P < 0.05$). For (**C**), the Mantel-Cox test was used, asterisks indicate $P$ <0.0001(****), "ns" denotes no significant difference. For (**D**), the independent samples t test was used, asterisks indicate $P$ <0.05(*), <0.001(***), "ns" denotes no significant difference. $P < 0.05$ was considered statistically significant.

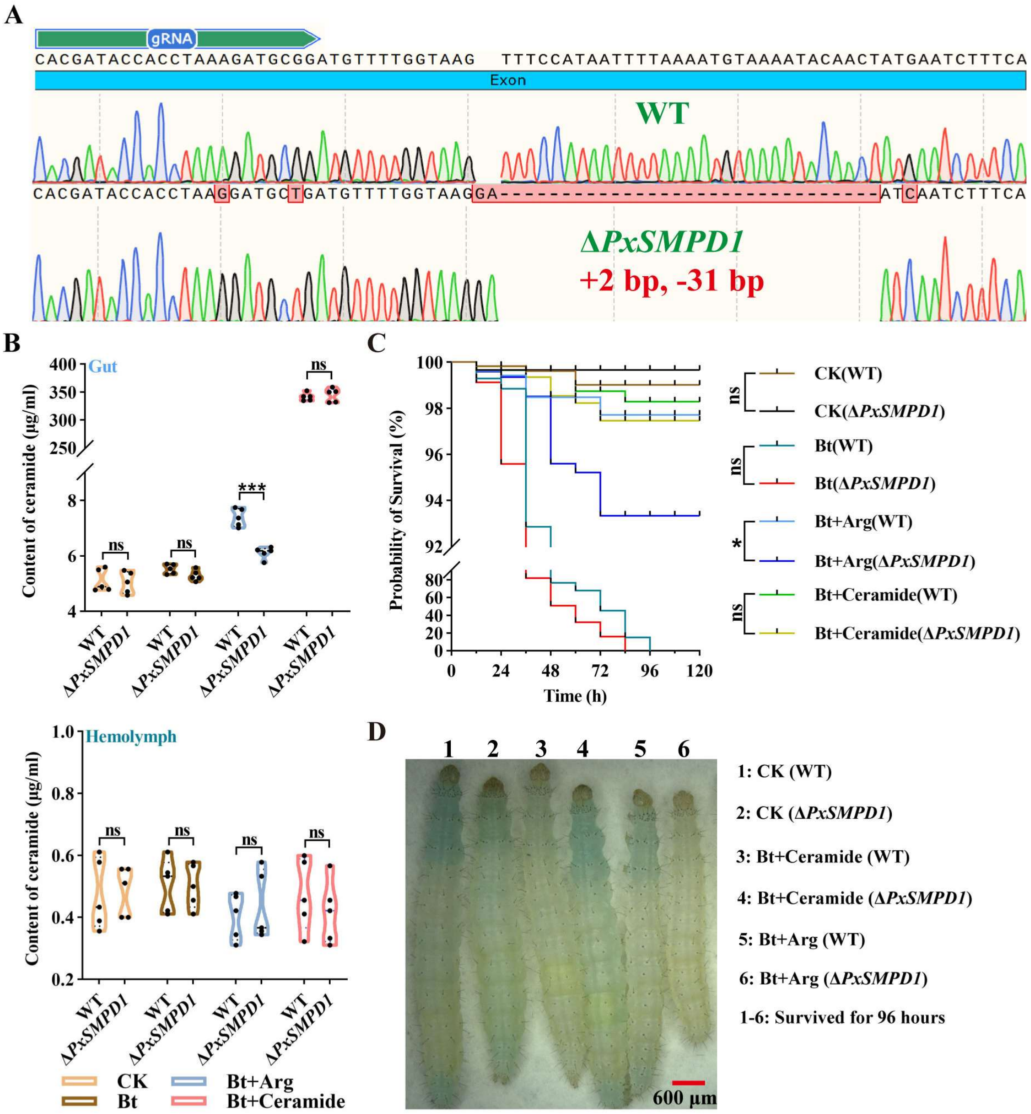


**S3 Fig. The ceramide synthesis pathway gene *PxSMPD1* mediates defense against Bt in *P. xylostella.*** (**A**) Generation of the mutant Δ*PxSMPD1* in *P. xylostella*; the mutant carries a +2 bp and –31 bp insertion / deletion (indicated as +2, –31 bp). (**B**) Deletion of *PxSMPD1* significantly attenuates the arginine induced increase in gut ceramide levels in *P. xylostella*; n = 5 biological replicates. (**C**) Deletion of *PxSMPD1* significantly reduces the ability of arginine to mediate defense against Bt in *P. xylostella*; n = 3 biological replicates. (**D**) Morphological characteristics of larvae from the treatment groups corresponding to (**C**) at 96 hours of growth. WT, wild type *P. xylostella*; Δ*PxSMPD1*, the mutant carrying a deletion of *PxSMPD1*. Abbreviations of the full names are used, with the details provided in the Materials and Methods section. For (**B**), the independent samples t test was used; asterisks indicate $P < 0.001$ (**), "ns" denotes no significant difference. For (**C**), the Mantel-Cox test was used, asterisks indicate $P < 0.05$(*), "ns" denotes no significant difference. $P < 0.05$ is considered statistically significant.

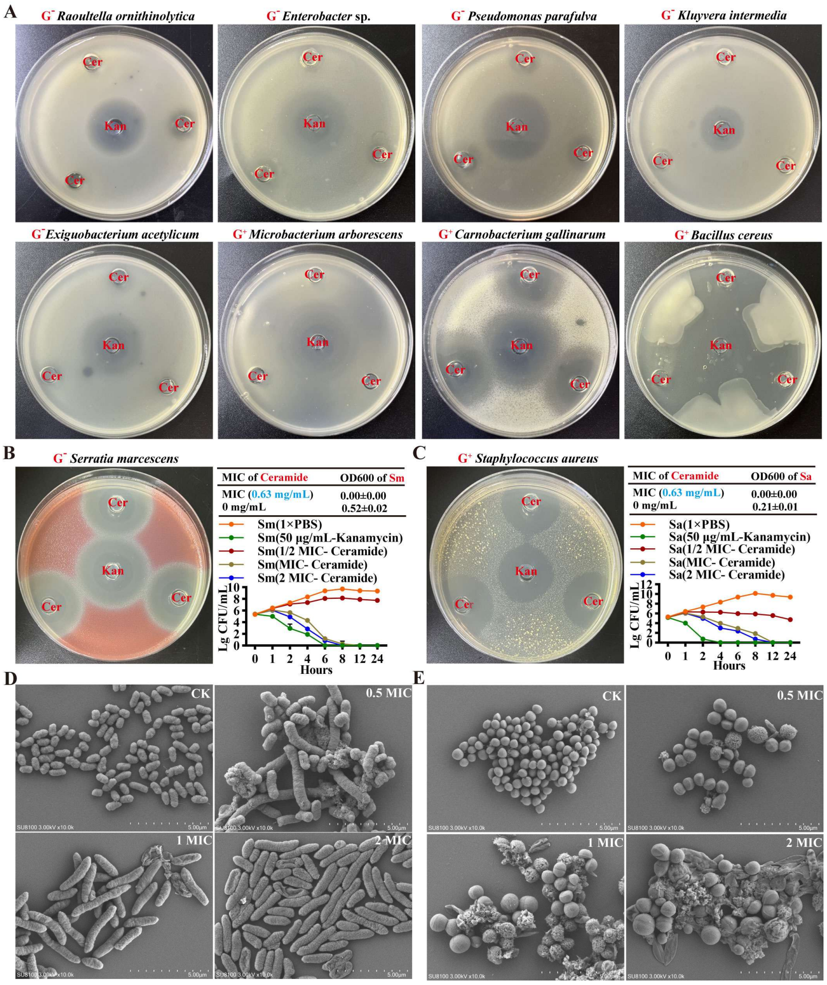


**S4 Fig. Mechanistic specificity of the antibacterial activity of ceramide.** (**A**) Ceramide exhibits broad-spectrum antibacterial activity against Gram-positive bacteria as shown by the Oxford cup inhibition zone method [51] on agar plates. (**B**) In *vitro* inhibition of *Serratia marcescens* (Sm) by ceramide: the plate assay shows the results of the Oxford cup inhibition zone method [51]; the three-line table presents the minimum inhibitory concentration (MIC) of ceramide against Sm; the curve graph displays the inhibition kinetics of Sm at different MICs of ceramide; n = 3 technical replicates. (**C**) In *vitro* inhibition of *Staphylococcus aureus* (Sa) by ceramide: the plate assay shows the results of the Oxford cup inhibition zone method [51]; the three-line table presents the MIC of ceramide against Sa; the curve graph displays the inhibition kinetics of Sa at different MICs of ceramide; n = 3 technical replicates. (**D**) Microscopic morphology of Sm treated with different MICs of ceramide. (**E**) Microscopic morphology of Sa treated with different MICs of ceramide. $G^-$ and $G^+$ denote Gram-negative and Gram-positive bacteria, respectively. Cer, ceramide; Kna, kanamycin; Sm, *Serratia marcescens*; Sa, *Staphylococcus aureus*; CK, control without ceramide treatment; 0.5 MIC, 1 MIC, and 2 MIC represent different concentrations of ceramide relative to the MIC.

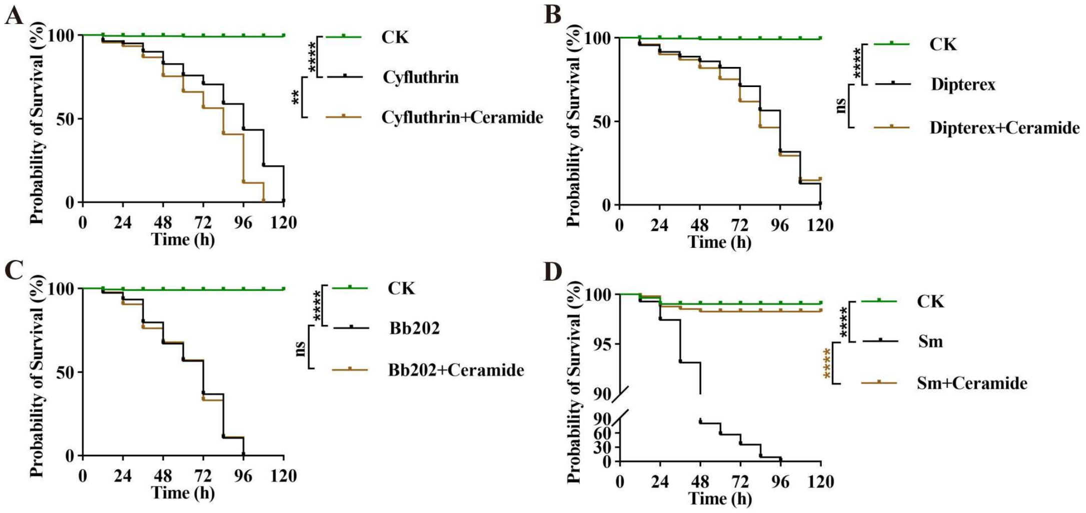


**S5 Fig. Ceramide specifically mediates defense against pathogenic bacteria in *P. xylostella.*** (**A**) Ceramide does not mediate defense against Cyfluthrin in *P. xylostella*; n = 3 biological replicates. (**B**) Ceramide does not mediate defense against Dipterex in *P. xylostella*; n = 3 biological replicates. (**C**) Ceramide does not mediate defense against Bb202 in *P. xylostella*; n = 3 biological replicates. (**D**) Ceramide mediates defense against Sm in *P. xylostella*; n = 3 biological replicates. Abbreviations of the full names are used, with the details are provided in the Materials and Methods section. For (**A**) to (**D**), the Mantel-Cox test was used, asterisks indicate $P < 0.01$ (**), < 0.0001 (****), "ns" denotes no significant difference. $P < 0.05$ is considered statistically significant.

**A**

**Presence of key ceramide synthesis enzymes in sphingolipid metabolism**

| Species | SPT | KDSR | CERS | SMPD | DES | GBA |
|---|---|---|---|---|---|---|
| ***Plutella xylostella*** | √ | √ | √ | √ | √ | √ |
| ***Drosophila melanogaster*** | √ | √ | √ | √ | √ | √ |
| ***Mus musculus*** | √ | √ | √ | √ | √ | √ |
| ***Danio rerio*** | √ | √ | √ | √ | √ | √ |
| ***Homo sapiens*** | √ | √ | √ | √ | √ | √ |
| ***Sus scrofa*** | √ | √ | √ | √ | √ | √ |
| ***Gallus gallus*** | √ | √ | √ | √ | √ | √ |

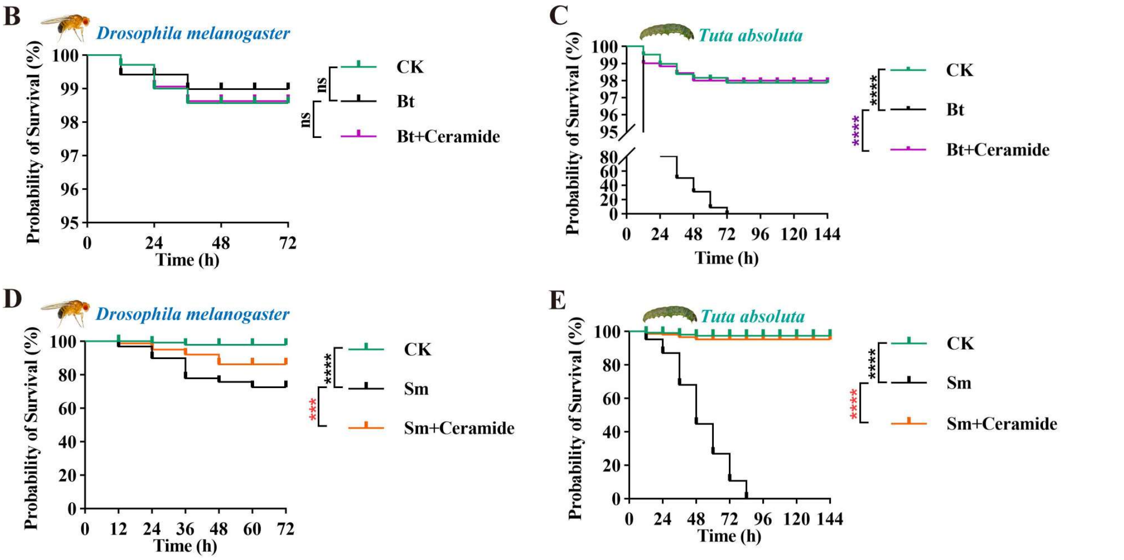


**S6 Fig. The antipathogenic effect of ceramide exhibits potential universality across insect species.** (**A**) Key enzymes of the ceramide synthesis pathway are widely present in eukaryotes. (**B**) *Drosophila melanogaster* does not exhibit sensitivity to Bt; n = 3 biological replicates. (**C**) Ceramide mediates defense against Bt in *Tuta absoluta*; n = 3 biological replicates. (**D**) Ceramide mediates defense against Sm in *Drosophila melanogaster*; n = 3 biological replicates. (**E**) Ceramide mediates defense against Sm in *Tuta absoluta*; n = 3 biological replicates. Abbreviations of the full names are used, with the details provided in the Materials and Methods section. SPT, KDSR, CERS, DES, SMPD, and GBA are the abbreviations for serine palmitoyltransferase, 3-ketodihydrosphingosine reductase, ceramide synthase, dihydroceramide desaturase, sphingomyelin phosphodiesterase, and glucocerebrosidase, respectively. Check marks "√" indicate presence in the NCBI database (National Center for Biotechnology Information). For (**B**) to (**E**), the Mantel-Cox test was used, asterisks indicate $P < 0.001$(***)、<0.0001(****), "ns" denotes no significant difference. $P < 0.05$ is considered statistically significant.

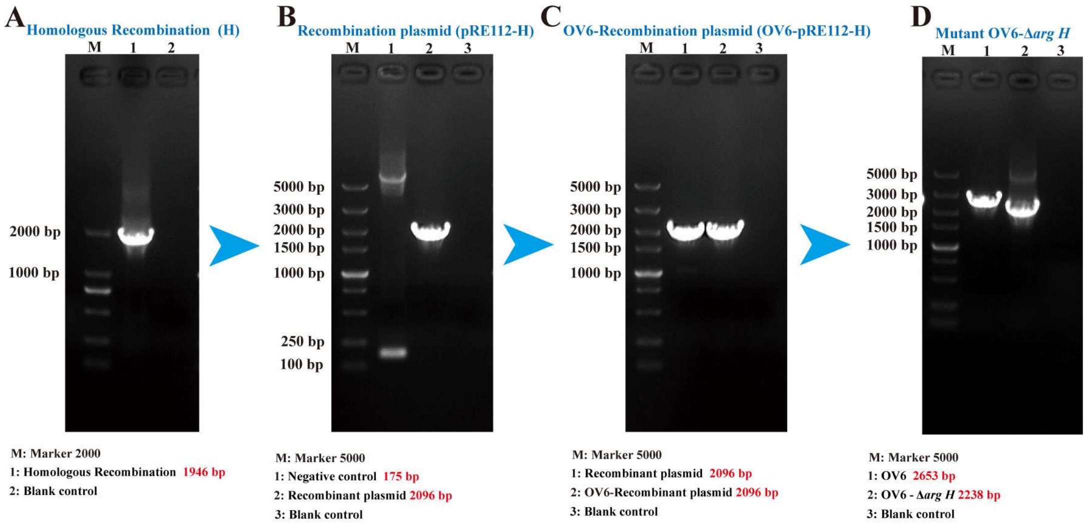


**S7 Fig. Construction of the OV6-Δ*argH* mutant strain.** (**A**) Generation of the homologous recombination fragment (H). (**B**) Introduction of the homologous recombination fragment (H) into the target plasmid (pRE112) to generate the recombinant plasmid (pRE112-H). (**C**) Transfer of the recombinant plasmid into the OV6 wild-type strain to generate the recombinant strain (OV6-pRE112-H). (**D**) Screening and selection of the OV6-Δ*argH* mutant strain carrying a deletion of the *argH* gene.

**S1 Table. Mobile phase elution gradient**

| Serial Number | Time（min） | %A | %B | %C | %D |
|---|---|---|---|---|---|
| 1 | 0.00 | 2.0 | 0.0 | 98.0 | 0.0 |
| 2 | 0.3 | 2.0 | 0.0 | 98.0 | 0.0 |
| 3 | 5.5 | 9.0 | 80.0 | 11.0 | 0.0 |
| 4 | 7.0 | 8.0 | 16.0 | 58.0 | 18.0 |
| 5 | 7.30 | 8.0 | 16.0 | 58.0 | 18.0 |
| 6 | 7.7 | 8.0 | 0.0 | 71.0 | 21.0 |
| 7 | 8.0 | 4.0 | 0.0 | 36.0 | 60.0 |
| 8 | 9.0 | 4.0 | 0.0 | 36.0 | 60.0 |
| 9 | 10.0 | 2.0 | 0.0 | 98.0 | 0.0 |
| 10 | 12.0 | 2.0 | 0.0 | 98.0 | 0.0 |

**S2 Table. Amplification and detection primers of OV6**

| Name | Primer sequence (5'→3') |
|---|---|
| ArgH-F | ATGGCACTTTGGGGCGGGCGTTTTA |
| ArgH-R | TTACACCAGACGCTGTTTCGCGT |
| Up-F | ACTTCGCGTAACACGGTACG |
| Up-R | CGCCGCTCCTGAATGATTCCA |
| Down-F | CTTCGCTAACCCGACCTACAG |
| Down-R | CTGATTAACACCCTGCTGGCC |
| KnaR-F | GGAATCATTCAGGAGCGGCGGACAGAGGTAGCTTGCAGTGG |
| KnaR-R | GTAGGTCGGGTTAGCGAAGGTCAGAAGAACTCGTCAAGAAGGCG |
| XmaI-F | cccccgggACTTCGCGTAACACGGTACG |
| XbaI-R | tgctctagaCTGATTAACACCCTGCTGGCC |
| Test1-F | ACTTCGCGTAACACGGTACG |
| Test1-R | CTGATTAACACCCTGCTGGCC |
| Test2-F | CATCCAACGCCATTCATGGC |
| Test2-R | CTTAAACGCCTGGtTGCTACGC |
| Test3-F | GCAGGTACATTTCCAGCTTGGG |
| Test3-R | CGAATAAGACGCTGCTGGCAT |

Note: The red bases are the homologous sequence of the R end of the upstream homology arm of the *ArgH* gene; the blue bases are the homologous sequence of the F end of the downstream homology arm of the *ArgH* gene; the lowercase letters represent the restriction enzyme sites.

**S3 Table. Amplification primers of *P. xylostella***

| Gene name | | Primer sequence (5’→3’) |
|---|---|---|
| | *Px017172*-F | AGCACAGCTACCTACGAAAC |
| | *Px017172*-R | TCAAAGAGCTTGCAATTCTGG |
| | *Px016398*-F | CCTCGGCTATCTGAACCAGC |
| | *Px016398*-R | CTTGCCCGTGAACTGGAAGC |
| | *Px017276*-F | CACAGAACACCTGAATGTGCCTCG |
| | *Px017276*-R | CCTTACCCAAGCCAGAAAGTCG |
| | *Px010122*-F | GGAATCGGCAAAGCCAGTG |
| | *Px010122*-R | CATACCCAGATGTGATGTCCAG |
| | *Px008942*-F | CTAAATACGCCAACTACCAGCGGT |
| | *Px008942*-R | GCTCGGACTGATGTGCTCCAT |
| | *Px008081*-F | GGCTGCTTCTCCAACCATTCTTC |
| Ceramide | *Px008081*-R | ACTTCACGACTAACGCGTCG |
| Biosynthesis | *Px004605*-F | CAAGGATCACGCTCGCAGAGAT |
| | *Px004605*-R | GTCTGTCGCTCGACTTCAGTGAT |
| | *Px003742*-F | CGATCTGGTGGATGTACGCAGTCT |
| | *Px003742*-R | CAGCTGTTCGTCGGTAGAACCGT |
| | *Px007275*-F | CAACTCTGTCTCGCAGAACCT |
| | *Px007275*-R | GTTTCGTCGTGGTAATACGGCT |
| | *Px009716*-F | CTGTCCAGTGCTCAGATCGT |
| | *Px009716*-R | ACGTACCAGCTTGCAGACCAG |
| | *Px009600*-F | CTACGAGTACATCACCGTCCCT |
| | *Px009600*-R | CGTTGAGGTTCCATTCCCAACT |
| | *Px007446*-F | GTTCTGGCTGCGCCTCTACTCAT |
| | *Px007446*-R | CCACACTGAGATCTTCGTCTGGT |
| | *PxLyszomes2*-F | AGTTGATAACTGACGACATCACGAA |
| | *PxLyszomes2*-R | GGCACTTGTTCTTCCATCCATAC |
| | *PxGloverin2*-F | GCCGGGCTATTCTTACAAGC |
| | *PxGloverin2*-R | CCCGTAGTCTGGTCTCCCAT |
| AMPs | *PxMoricin3*-F | GAAGGGCGGCCATGTCATTA |
| | *PxMoricin3*-R | TACCCCTGGTTCCTGTTCCG |
| | *PxCecropin2*-F | TTCGTGTTCGTGGCTGTTTT |
| | *PxCecropin2*-R | GGCCAACTCTTTCCAATTTCTT |
| Internal | *RPL32*-F | CAATCAGGCCAATTTACCGC |
| reference | *RPL32*-R | CTGGGTTTACGCCAGTTACG |

**S4 Table. Amplification and detection primers of *P. xylostella*.**

| Name | Primer sequence (5’→3’) |
|---|---|
| *Px003742*-F | ATGAAAGAGCTGTTGTATTTGTTTACGGTGC |
| *Px003742*-R | TTAACTGAAAAACAGGTCTCTCAGTTGTTG |
| *Px008081*-F | ATGGGTGCTAAAGTATCCCGTAC |
| *Px008081*-R | TTAGCTCTCCAGCCCTCTATG |
| M13-F | TGTAAAACGACGGCCAGT |
| M13-R | CAGGAAACAGCTATGACC |
| *Px003742*-gRNA-F | TAATACGACTCACTATAGG<u>CACGATACCACCTAAAGATG</u>GT<br>TTTAGAGCTAGAAATAGCAAGTTAAAATAAGGCTAGTCC |
| *Px008081*-gRNA-F | TAATACGACTCACTATAGG<u>CCAAACAGTCGGTTGTGGAG</u>G<br>TTTTAGAGCTAGAAATAGCAAGTTAAAATAAGGCTAGTCC |
| gRNA-comR | AAAAGCACCGACTCGGTGCCACTTTTTCAAGTTGATAA<br>CGGACTAGCCTTATTTTAACTTGCTATTTCTAGCTCTAAAA |
| *Px003742*-T-F | ACCTAGAATGCCTGATCTGCC |
| *Px003742*-T-R | GCGTCTGTTAGTACTGCGAT |
| *Px008081*-T-F | TGTCCACCGATATCTTCGC |
| *Px008081*-T-R | CAGCTGAATCACCAGATTAACC |

Note: The red bases represent the T7 promoter sequence; the underlined bases represent the sgRNA recognition sequence; the blue bases represent the sgRNA scaffold sequence.
